# Meta-organization of Translation Centers Revealed by Proximity Mapping of Endoplasmic Reticulum Ribosome Interactors

**DOI:** 10.1101/398669

**Authors:** Alyson M. Hoffman, Christopher V. Nicchitta

## Abstract

The endoplasmic reticulum (ER) is a nexus for mRNA localization and translation; the molecular organization of these processes remains however largely undefined. To gain insight into mechanisms supporting a diverse ER translational landscape, we utilized BioID labeling to study the protein neighborhoods of the Sec61 translocon, specifically Sec61β, an established ribosome interactor, and ER proteins (Ribophorin I, LRRC59, and Sec62) previously implicated in ribosome association. Divergent protein interactomes enriched for distinct GO functions were identified for the four reporters, within a cohort of shared interactors. Efficient BioID tagging of ribosomes was only observed for the Sec61β and LRRC59 reporters. RNA-seq analyses of the Sec61β- and LRRC59-labeled ribosomes revealed divergent enrichments in mRNAs and identified a transcriptome-wide role for the ER in proteome expression. These data provide evidence for a mesoscale organization of the ER and suggest that such organization provides a mechanism for the diversity of translation on the ER.

## Introduction

The endoplasmic reticulum (ER) is a highly heterogeneous organelle composed of both rough and smooth membrane domains, distinguished by the presence or absence of bound ribosomes, a contiguous sheet and tubular membrane organization, and a diversity of primary cellular functions including secretory/membrane protein biogenesis, lipid biosynthesis, and calcium storage (English & Voeltz, 2013; Fawcett, 1966; Lynes & Simmen, 2011; Schwarz & Blower, 2016). In addition to these defining features, the ER engages in membrane-membrane communication with different organelles, including mitochondria, endosomes, and the plasma membrane (English & Voeltz, 2013; Helle et al., 2013; Valm et al., 2017). The sites of organelle–ER communication are marked by multi-protein assemblies that define areas of regional specialization, e.g., mitochondrial-associated membranes (MAMs), and provide evidence for the spatial organization of the membrane proteome as a biochemical/biophysical mechanism to accommodate the various structural and functional properties of this critical organelle (de Brito & Scorrano, 2010; Hung et al., 2017; Jing et al., 2015; Vance, 2014).

In addition to a dedicated function in secretory/membrane protein biogenesis, recent and past studies examining the subcellular distribution of mRNAs between the cytosol and ER compartments have revealed a transcriptome-wide role for the ER in proteome expression (Mueckler and Pitot 1981; Diehn et al. 2000; Diehn et al. 2006; Stephens and Nicchitta 2008; Jan, Williams, and Weissman 2014; Chartron, Hunt, and Frydman 2016; Voigt et al. 2017; Reid and Nicchitta 2012). The high enrichment in ER localization and translation of secretory/membrane protein-encoding mRNAs further validates the established function of the ER in secretory and membrane protein biogenesis (Reid and Nicchitta 2012; Jan, Williams, and Weissman 2014; Chartron, Hunt, and Frydman 2016). An unexpected yet consistent finding in these studies is the comparatively modest enrichment of cytosolic mRNAs on free, cytosolic ribosomes (Reid and Nicchitta 2012; Jan, Williams, and Weissman 2014; Chartron, Hunt, and Frydman 2016). Given that eukaryotic transcriptomes are substantially weighted to cytosolic protein-encoding mRNAs, such modest enrichments indicate a broad representation of these mRNAs on the ER and suggest a global role for the ER in transcriptome expression (Diehn et al., 2006; Mueckler & Pitot, 1981; Reid & Nicchitta, 2015; Voigt et al., 2017). Although a function for the ER in the translation of cytosolic protein-encoding mRNAs has been under investigation for many decades, more recent biochemical and structural biology studies of the ribosome-Sec61 translocon, composed of Sec61α, Sec61β and Sec61γ subunits, would suggest that such a function is unlikely (Becker et al., 2009; Pfeffer et al., 2014, 2015; Prinz, Behrens, Rapoport, Hartmann, & Kalies, 2000; Schaletzky & Rapoport, 2006; Voorhees, Fernández, Scheres, & Hegde, 2014). The Sec61 translocon serves as a ribosome receptor and translocation channel, where the ribosomal protein exit channel resides in close physical apposition to the translocon channel entrance site. Thus, Sec61 translocon-associated ribosomes are presumed to be dedicated to the translation of secretory and membrane protein-encoding mRNAs. Recent work supports additional mechanisms of supporting ER-localized translation, where translation of cytosolic protein-encoding mRNAs on the ER could be accommodated by Sec61 translocon-independent ribosome association mechanisms (Cui, Zhang, and Palazzo 2012; Voigt et al. 2017; Reid and Nicchitta 2012; Stephens and Nicchitta 2008; Potter and Nicchitta 2000). In support of this view, a number of ER resident membrane proteins other than the Sec61 translocon have been proposed to function as ribosome receptors, including the oligosacharyltransferase (OST) subunit Ribophorin I, LRRC59 (p34), and p180 (RRBP1) (Harada, Li, Li, & Lennarz, 2009; Kreibich, Freienstein, Pereyra, Ulrich, & Sabatini, 1978; Savitz & Meyer, 1990; Tazawa et al., 1991). Although these proposed ribosome receptors have been shown to display high binding affinity to ribosomes *in vitro*, very little is known regarding the potential diversity of ribosome-ER protein interactions *in vivo*.

Here we utilized BioID *in vivo* proximity mapping as a means to investigate the interactomes of a subset of proposed ribosome receptors. These data revealed a higher order (mesoscale) organization of the ER, where the BioID reporters reside in stable protein environments with clear GO functional enrichments, consistent with the organization of the ER membrane into discrete translation centers. Of the four candidates examined, only two, Sec61β and LRRC59, efficiently labeled ER-bound ribosomes. Intriguingly, RNA-seq analysis of ribosomes from the two distinct sites revealed both enriched and shared transcriptome cohorts. These data are consistent with a higher order organization of the translation functions of the ER into translation centers, which we define as enriched assemblies of interacting proteins, associated ribosomes, and bound mRNAs. In addition, these data suggest mechanisms whereby the ER could serve a broad role in proteome expression.

## Results

### Experimental Overview

Recent reports identify a transcriptome-wide function for ER-associated ribosomes in proteome expression (Reid and Nicchitta 2012; Voigt et al. 2017; Jan, Williams, and Weissman 2014; Chartron, Hunt, and Frydman 2016). Given the central role of the Sec61 translocon as both the protein-conducting channel and ribosome receptor for membrane and secretory proteins (Gorlich, Prehn, Hartmann, Kalies, & Rapoport, 1992; Voorhees et al., 2014), we postulated that ER membrane proteins other than the Sec61 translocon participate in ribosome-ER interactions, as a mechanism to support transcriptome expression on the ER (Pfeffer et al., 2015; Reid & Nicchitta, 2015; Voorhees et al., 2014). To test this hypothesis, we used a BioID proximity labeling approach, where BioID chimera of known or proposed ribosome interacting proteins were used to map both ER protein-ribosome and proximal ER membrane protein interactions (**Figure 1A,B**). Briefly, BioID uses a mutant bacterial biotin ligase (BirA*) that releases an unstable, amine-reactive biotin intermediate (biotin-AMP) from the active site; biotin-AMP can then react with near-neighbor proteins and thus provide *in vivo* protein-protein spatial interaction information (Roux, Kim, Raida, & Burke, 2012). As schematically illustrated in **Figure 1A**, ER membrane protein-BioID chimera constructs were engineered, inducible HEK293 Flp-In cell lines generated, and interactomes studied by biotin addition followed by subcellular fractionation and proteomic analyses of biotin-tagged proteins. As noted above, prior studies have established the Sec61 translocon as a ribosome receptor and so Sec61β-BioID was selected to report on the Sec61 translocon interactome (Deshaies, Sanders, Feldheim, & Schekman, 1991; Levy, Wiedmann, & Kreibich, 2001; Pfeffer et al., 2015). Ribophorin I, a component of the OST, is vicinal to translocating nascent chains, has been proposed to function as a ribosome receptor, and was found to be a Sec61 translocon interactor, thus complementing the Sec61β-BioID interactome screen (Harada et al., 2009; Kreibich et al., 1978; Wild et al., 2018). Sec62, though relatively unstudied in mammalian systems, is orthologous to yeast Sec62, which participates in post-translational translocation, and in mammalian systems has been demonstrated to function in ribosome binding, with binding interactions mapped to regions adjacent to the ribosome exit tunnel (Lang et al., 2012; Müller et al., 2010). LRRC59, also relatively unstudied, was identified as a ribosome binding protein through biochemical reconstitution approaches and chemical crosslinking, where it was demonstrated to reside near large ribosomal subunits (Ichimura et al., 1993; Tazawa et al., 1991). An *in vivo* function for LRRC59 in ribosome binding has not been established, although anti-LRRC59 IgG and Fab inhibit ribosome binding and protein translocation *in vitro* (Ichimura et al., 1993). These four proteins were chosen as belonging to both well-studied complexes, e.g., Sec61β and Ribophorin I, and little studied proteins lacking a defined function in mammalian cells, e.g., LRRC59 and Sec62, to provide an expanded understanding of the molecular organization of the ER.

**Figure 1.**
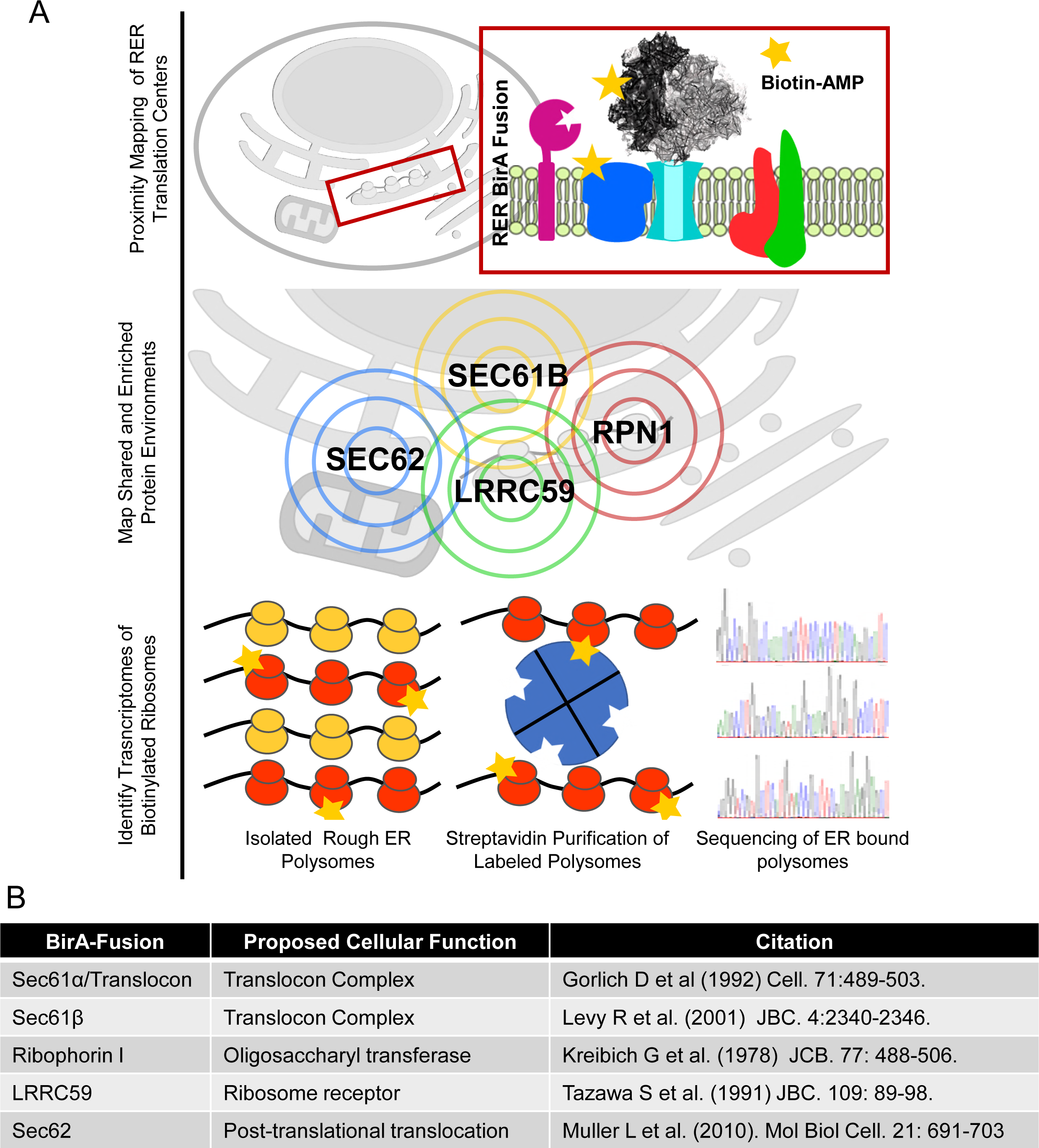
Experimental approach to the analysis of ribosome interactors and localized translation on the endoplasmic reticulum (ER). **A)** Schematic of experimental goals. Stable cell lines expressing inducible BirA* fusion proteins of previously identified ribosome associated membrane proteins, listed in (**B**), were prepared and used to determine the near-neighbor protein interactomes for each via the BioID method. In addition to determining candidate ribosome interactor protein neighborhoods, reporter-mediated ribosome labeling was examined. To define the transcriptomes present at each of the postulated ER translation centers, biotin-tagged, ER-associated ribosome were recovered by detergent solubilization and avidin affinity isolation. Transcriptome compositions were determined by RNA-seq. **B)** Summary listing of candidate ribosome-interacting proteins, proposed functions, and linked citation.

### Ribosome Interactor-BioID Chimera are ER-Localized and Predominately Label ER Membrane Proteins

To assess the subcellular localization and proximity labeling activity of the ribosome interactor-BioID chimera introduced above, reporter cell lines were induced for 16 hours with biotin supplementation and the subcellular reporter expression patterns determined by immunofluorescence, using antisera directed against BirA, with biotin labeling patterns examined by staining with a streptavidin-AF647conjugate (**Figure 2A**). A cell line containing the cloning vector backbone served as a negative control in this analysis (empty vector). As depicted in **Figure 2 A and B**, all four reporter cell lines displayed clear perinuclear reticular staining with the BirA antisera, consistent with an ER localization for all BioID chimera. Ribophorin I and Sec61β are subunits of oligomeric protein complexes; to ensure correct localization of these chimeras we also compared the hydrodynamic behavior of the BirA* chimera with the respective natively expressed proteins in the engineered BioID and empty vector cell lines by glycerol gradient velocity sedimentation (Nikonov, Snapp, Lippincott-Schwartz, & Kreibich, 2002)(**Figure 2 – figure supplement 1**). The sedimentation patterns of all BioID reporters and respective native proteins were similar to those of the empty vector-engineered cell lines and with sedimentation velocities consistent with their estimated complex sizes, suggesting that the chimera were appropriately assembled into native complexes (Harada et al., 2009). In yeast, such chimera complement genomic deletions of the parent gene, also indicative of native-like function (Jan, Williams, and Weissman 2014).

**Figure 2.**
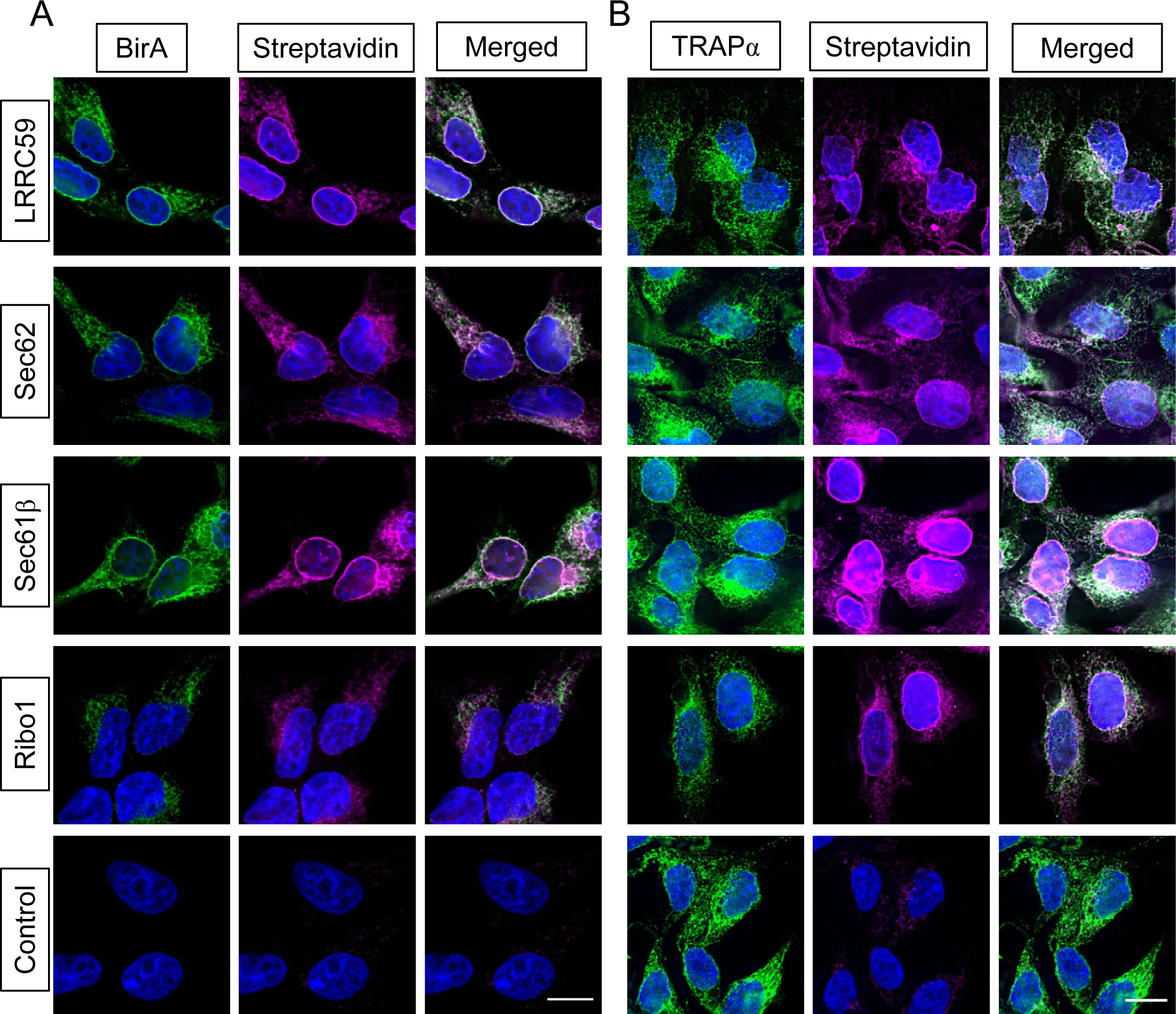
BioID reporters display ER-restricted subcellular localization and biotin-labeling activity. **A)** Immunofluorescence micrographs of each reporter cell line after 24 hours of doxycycline-induced expression (BirA channel) and overnight biotin treatment (streptavidin channel). The merged images reveal high coincidence of ER membrane figures and proximity labeling. Scale bar = 10 μm**. B)** Immunofluorescence micrographs of each reporter cell line showing colocalization of the resident ER membrane protein marker (TRAPα) and the biotin labeling pattern. Scale bar = 10 μm. Data shown are representative of two biological replicates.

Intriguingly, streptavidin staining of proximal biotin labeled targets co-localized with the BirA staining patterns, suggesting that the primary interactomes are largely confined to the ER (**Figure 2A,B**). These findings were further validated in studies comparing the streptavidin staining patterns with the resident ER membrane protein TRAP*α* (**Figure 2B**). As with the data depicted in **Figure 2A**, we observed extensive overlap of streptavidin staining pattern with that of the resident ER protein marker. Surprisingly, after 16 h of induction/biotin labeling the biotin tagging pattern was highly restricted to the ER, with little discernible tagging of cytosolic proteins.

The immunofluorescence data (**Figure 2**) were further evaluated in cell fractionation studies of the BirA* chimera and the biotin labeling patterns (**Figure 3**). Using a previously validated sequential detergent fractionation protocol (Jagannathan, Nwosu, & Nicchitta, 2011; Stephens & Nicchitta, 2007), where the cytosol fraction is released upon addition of a digitonin-supplemented buffer and the membrane fraction subsequently recovered by addition of a NP40/sodium deoxycholate/high-salt buffer, BioID chimera distributions were assessed by SDS-PAGE/immunoblot analysis. These data are depicted in **Figure 3** and demonstrate that all ER membrane protein reporters were wholly recovered in the membrane fraction (**M**) and displayed SDS-PAGE mobilities consistent with their predicted molecular weights. Mirroring the immunofluorescence data shown in **Figure 2A and B**, proximal protein biotin labeling was highly enriched in the membrane fractions (**M**) (**Figure 3B,C**,TRAP*α* as ER marker), with only modest labeling of cytosolic proteins (**C**) (**Figure 3B,C**, β-tubulin as cytosolic marker). Interestingly, the biotin labeling patterns of the membrane fractions were readily distinguishable from one another, suggesting that the ER protein-reporters reside in distinct protein neighborhoods. The relatively paucity of biotinylated cytosolic proteins, visualized by both immunofluorescence staining and direct biochemical analysis, was unexpected. Because the reactive biotin-AMP intermediate diffuses from the BirA* active site to modify accessible lysine residues of proximal proteins (Choi-Rhee, Schulman, & Cronan, 2004), we expected that membrane and cytosolic proteins would be similarly accessible to modification. The bias to biotin-conjugation of ER membrane proteins suggests that the labeling radius for the reactive biotin-AMP intermediate is highly restricted (Kim et al. 2014).

**Figure 3.**
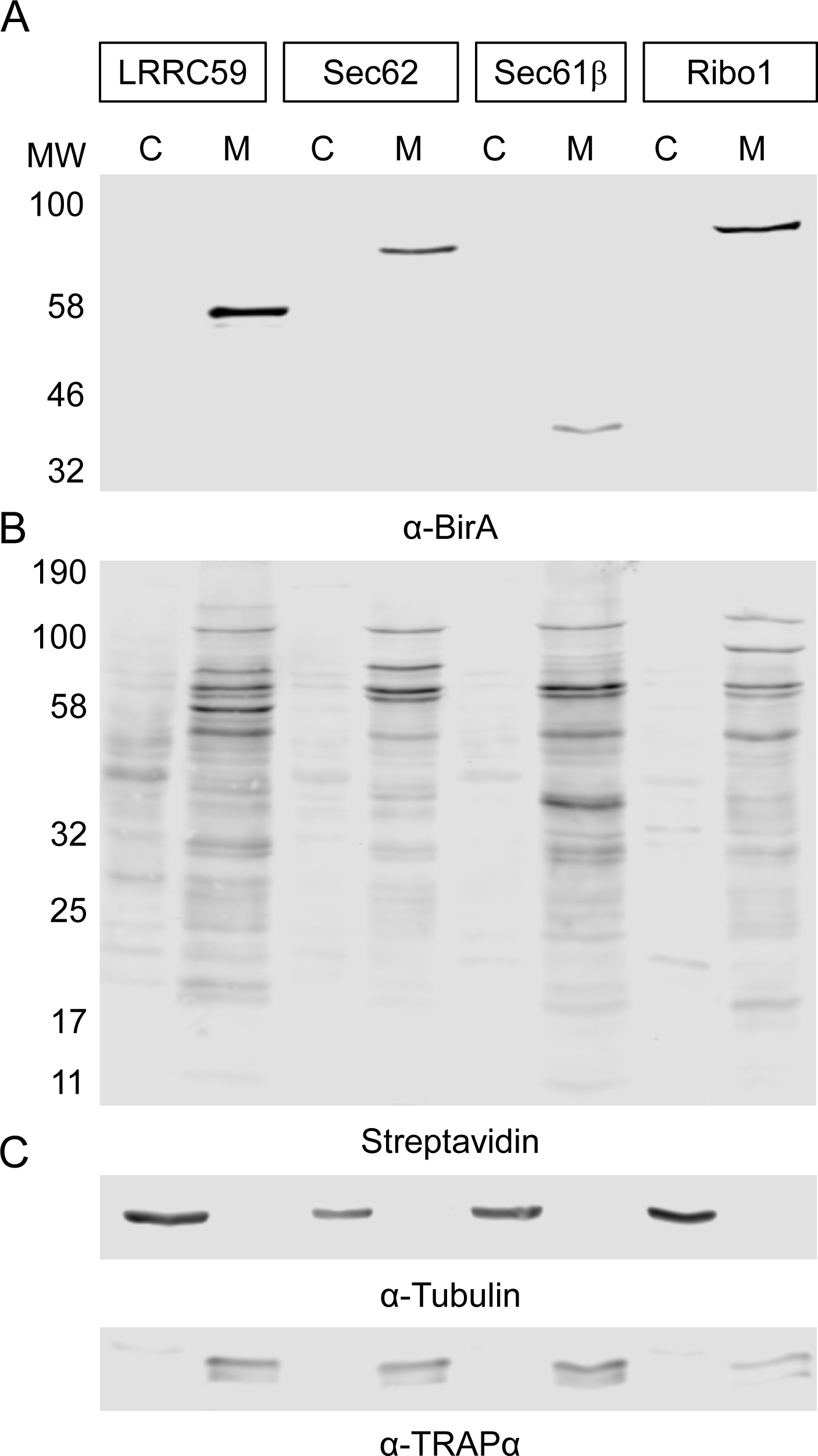
Biochemical fractionation of BioID reporter cell lines demonstrates localization of the reporter constructs to the membrane fraction and highly enriched biotin tagging of membrane proteins. **A)** BirA immunoblot depicting the localization of each construct to the membrane fraction of the detergent fractionated cells. BirA*-mediated biotin labeling, depicted by streptavidin blot, reveals distinct labeling patterns for each cell line and high enrichment of the ER fraction over the cytosolic fraction. **C)** Cytosol and ER marker protein distribution in the cytosol and membrane fractions derived from the four BioID reporter cell lines. Cytosol marker = β-tubulin; ER marker = TRAP*α*. Data shown are representative of two biological replicates.

### Evidence for meso-organization of ER membrane protein assemblies

BioID proximity labeling experiments are typically conducted over many hours (Varnaitė; & MacNeill, 2016) (e.g. 16-24h), a reflection of the slow release kinetics of the reactive biotin-AMP catalytic intermediate from the BirA* active site (Kwon & Beckett, 2000). Because this time scale is substantially slower than that of most cellular processes, the specificity of the labeling reaction is formally a concern (Rees, Li, Perrett, Lilley, & Jackson, 2015), though it has been demonstrated that neighboring proteins can be distinguished from random interactors by their higher relative labeling over non-specific controls (Rees et al. 2015; Kim et al. 2014; Roux et al. 2012; Ueda et al. 2015; Gupta et al. 2015). In the context of the experiments with ER membrane-localized BioID reporters, we were concerned that reporter diffusion in the constrained 2D environment of the ER membrane over such extended labeling times could confound identification of near-neighbor and interacting proteins. To address this concern, we examined the BioID reporter biotin labeling patterns as a function of labeling time. Our prediction was that the biotin labeling patterns would diversify as labeling times increased, a consequence of the expected random diffusion of the reporter chimera within the 2-D constraints of the ER membrane. The results of these experiments are shown in **Figure 4A**. Depicted are streptavidin blots of the cytosol and ER protein fractions from the four BioID reporter cell lines, sampled over a time course of 0–6 hours. Two observations are highlighted here. First, as noted above, the relatively enhanced labeling of membrane proteins (**M**) to cytosolic proteins (**C**) is evident throughout the time course examined, with very modest levels of cytosolic protein labeling throughout the time course for all constructs. Second, contrary to our expectations, the membrane protein biotin labeling patterns did not substantially diversify over the labeling time course (**Figure 4A**). Rather, the labeling pattern intensified as labeling time increased. The biotin labeling patterns revealed by SDS-PAGE were further analyzed by densitometric analysis (**Figure 4A**), where it can be appreciated that the biotin labeling patterns intensify, but only modestly diversify, as a function of labeling time. These data suggest that the BioID interactomes comprise largely stable membrane protein assemblies (**Figure 4B),** rather than the presumed randomizing interactomes expected of diffusion-based interactions (**Figure 4B**) (Goyette & Gaus, 2017; Kusumi et al., 2012; Kusumi, Suzuki, Kasai, Ritchie, & Fujiwara, 2011; Singer & Nicolson, 1972).

**Figure 4.**
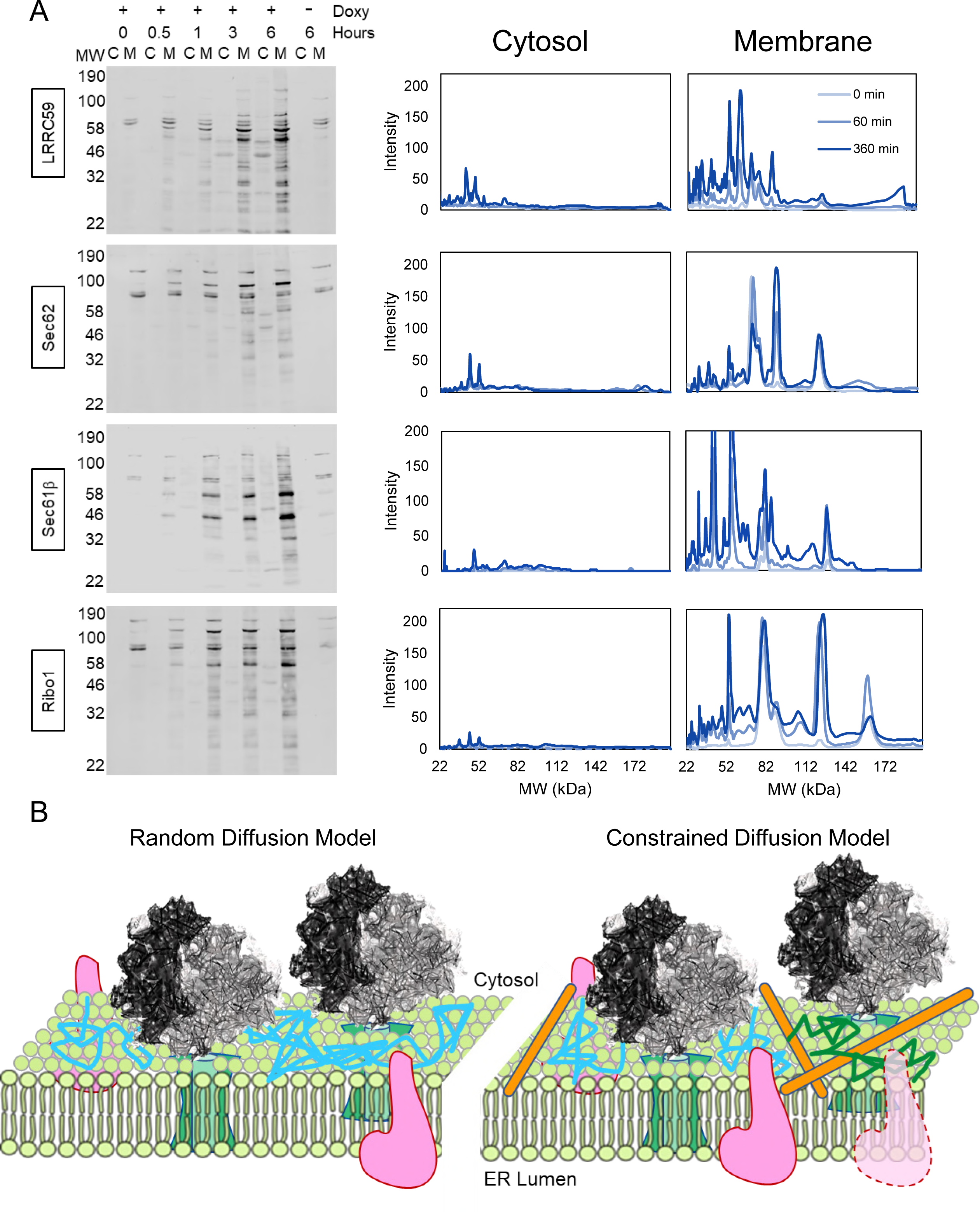
The BioID labeling patterns of the reporter cell lines largely intensify, rather than diversify, over time. **A)** Streptavidin blots of biotin labeling time courses are shown for each reporter cell lines. Also provided are line-intensity plots of selected time points through the six hour (hr) labeling period. Indicated are the biotin treatment time periods. **-Doxy** represents a six hr biotin treatment without prior doxycycline treatment, to test for leaky expression. **B)** Cartoons depicting the two predicted models of membrane protein diffusion. The leftward schematic (random diffusion) model depicts a biological membrane in which proteins diffuse freely in the 2-D membrane plane, encountering targets by random collision. The rightward schematic (constrained diffusion model), predicts that an organizing force, be it protein-protein interaction, lipid-enriched domains, or both, enables the formation of distinct compartments where protein diffusion is restricted. Data depicted in **A)** is representative of two biological replicates.

The data presented above (**Figure 4A**) are consistent with a model where the mobility of the BioID reporters is constrained (**Figure 4B**), perhaps reflecting mesoscale organization of the ER via biomolecular interactome networks, as has been extensively documented in the plasma membrane (Goyette & Gaus, 2017; Kusumi et al., 2012, 2011). We also considered that the labeling patterns could be influenced by either ER distribution biases (e.g., tubules vs. sheets) or protein-specific differences in reactivity to the biotin-AMP reactive intermediate. To examine these scenarios, we performed proximity labeling time course experiments using canine pancreas rough microsomes (RM), which lack the native topological structure of the ER, and a soluble, recombinant BirA*. Using this system, the reactive intermediate was delivered in *trans* and accessible to the ER surface by solution diffusion. The results of these experiments are shown in **Figure 5** and demonstrate that when accessible to RM proteins in *trans*, biotin labeling is pervasive and monotonic, with a diversity of proteins undergoing labeling and relative labeling intensities increasing as a function of labeling time (compare **Figure 5B** to **Figure 5D**). Combined, the data depicted in **Figures 4** and **5** indicate that these BirA* chimera are restricted to protein interactome domains of the ER *in vivo*, as represented by the distinct labeling patterns identified in each cell line.

**Figure 5.**
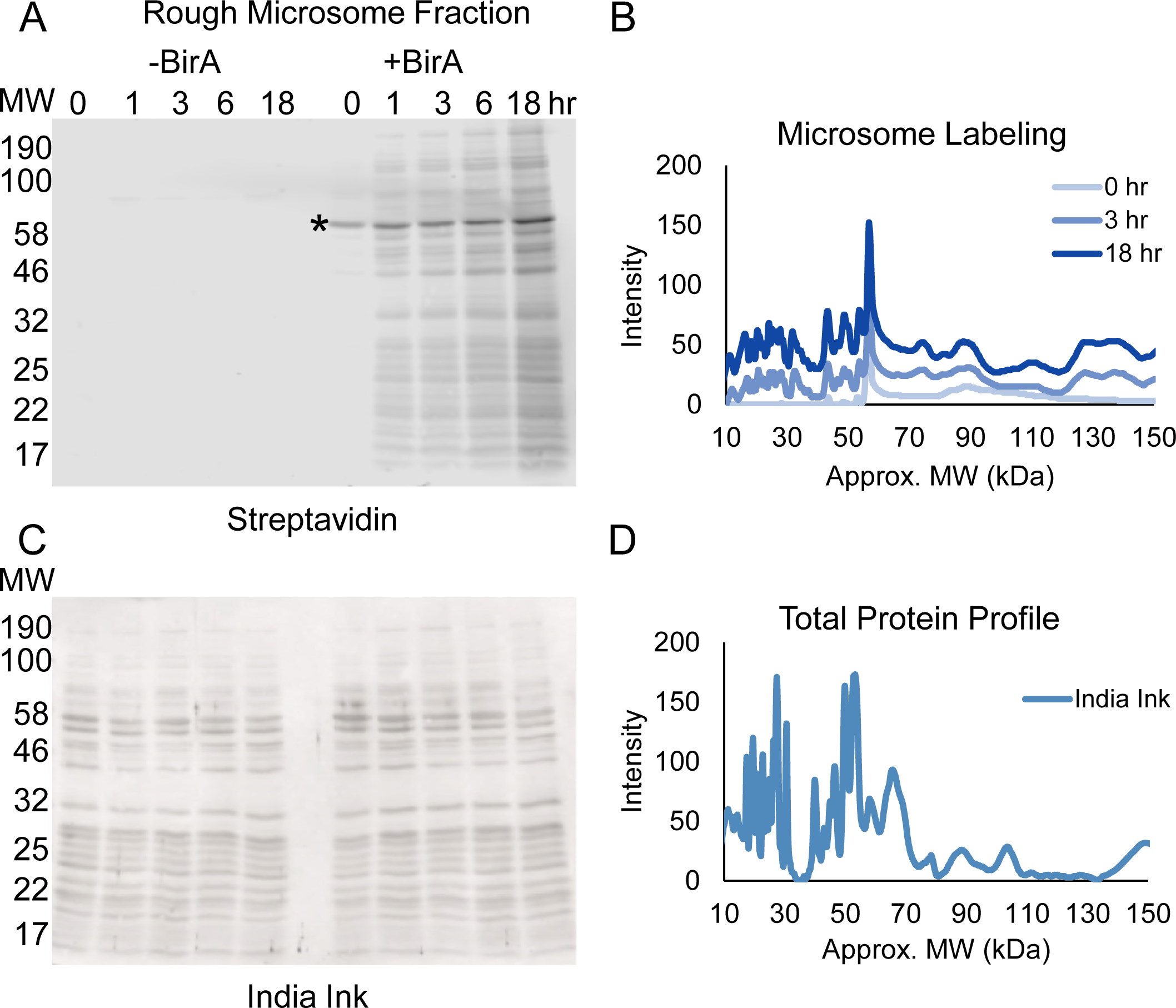
In the absence of *in situ*-like ER membrane organization and with *trans*-delivery of the reactive biotin intermediate, proximity-based selective labeling is abolished. **A)** SDS-PAGE gel depicting an *in vitro* labeling experiment conducted with canine rough microsomes (RM) incubated in the presence of soluble, recombinant BirA*. RM were incubated in the presence of an ATP regenerating system, biotin, and either BirA* or PBS as indicated. Note that the general avidin labeling pattern mirrors the total protein when the reporter is present in *trans*. Asterisk indicates the BirA*-GST fusion protein, which is biotinylated during its induction in *E.coli*. **B)** Lane intensity plots demonstrate a general increase at all molecular weights, indicating loss of specificity when the reporter is presented in *trans*. **C)** India ink stain of the blot above demonstrating equivalent protein loading for all samples. **D)** Lane intensity plot of the India ink stain, illustrating the overall similarity in the labeling of total accessible protein. Data shown are representative of two biological replicates.

To gain molecular insight into the protein neighborhoods of the reporters, cell lines were supplemented with biotin for 3 hours, the time point at which labeling intensity was highest compared to background, as illustrated in **Figure 4**, and biotinylated proteins captured from membrane extracts by streptavidin-magnetic bead affinity isolation. Elution was performed by biotin competition at high pH to select against non-specific background, with protein composition determined by mass spectrometry of the eluted samples. A summary of the analysis schema is depicted in **Figure 6A.** In brief, spectral counts of proteins meeting high confidence (1% FDR) cutoffs were normalized to those of natively biotinylated proteins and the subset with enrichments of > 2.5-fold over an empty vector control were selected. For these, two categories were defined; “enriched”, displaying an enrichment of > 2-fold over the combined normalized value, and “shared”, for those below this selection threshold. For all reporters, the majority of the labeled proteins meeting initial significance criteria were membrane proteins, corroborating the data presented in **Figures 2-4.** In the shared category, representing those proteins that met selection criteria and were present at similar normalized levels in two or more reporter datasets, about 80% were membrane proteins, 10% were cytoplasmic proteins, and 10% were nuclear proteins. Within the shared membrane protein category, a number of prominent ER resident membrane proteins were identified in the reporter datasets and included signal peptidase complex subunit 2, the eIF2*α* kinase PERK, DNAJC1, the ERAD-associated E3 ubiquitin ligase TRIM13, ERGIC-53, ER calcium ATPase 2, and reticulon-4. Other prominent ER resident proteins present in at least 3 of the 4 reporter datasets included Sec63 homolog, calnexin, and NADPH cytochrome P450. With respect to the enriched datasets, the candidate ribosome interactors LRRC59 and Ribophorin I returned identical numbers of neighboring/candidate interacting proteins (35), with Sec61β returning 19 enriched hits, and Sec62 11 enriched hits; these specific interactomes are discussed further below. In summary, proteomic analysis of the neighboring proteins for the indicated BioID chimera revealed a high enrichment in ER membrane proteins and within this category, proteins with established functions in canonical rough ER functions such as protein translocation/protein processing, the unfolded protein response (UPR), and ER-associated protein degradation (Cross, Sinning, Luirink, & High, 2009; Gardner, Pincus, Gotthardt, Gallagher, & Walter, 2013; Hayashi-Nishino et al., 2009; Vembar & Brodsky, 2008).

**Figure 6.**
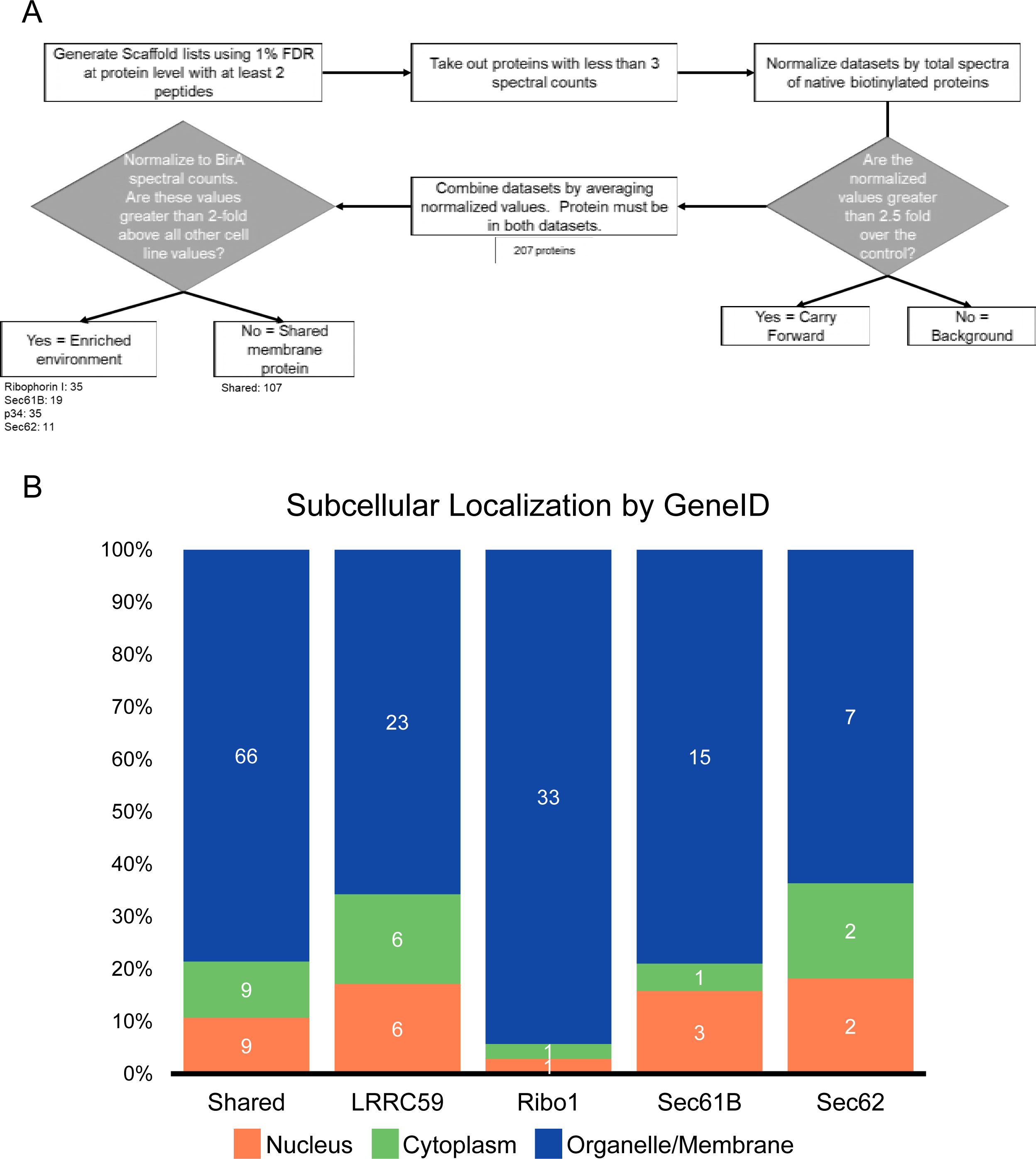
MS analysis of BioID-labeled proteins demonstrates a high labeling enrichment of membrane vs. cytoplasmic proteins. **A)** Schematic depicting the data analysis pipeline and significance selection criteria. All MS experiments were performed in biological duplicate. **B)** Stack plots depicting the relative distributions of cellular localization for enriched and shared proteins from each cell line. The number of genes in each category is embedded in each bar, to enable comparisons between reporter cell lines.

### Candidate Ribosome Interactors Reside In Distinct ER-Protein Neighborhoods

To further evaluate the subsets of neighboring proteins scoring in the enriched category, the datasets were visualized in Cytoscape (Shannon et al., 2003), cross-referenced via the STRING Protein-Protein Interaction Network resource (Szklarczyk et al., 2017), and functional enrichments for each reporter neighborhood determined by GO analysis. For clarity, the Cytoscape-generated plots depicted in **Figure 7** include the top enriched hits, with screened, shared interactors included as **Figure 8A-B**. The plots provided in **Figure 7** are coded to illustrate interactions with soluble proteins (yellow border) and membrane proteins (turquoise border). Centered and uncolored nodes indicate the chimera protein from each cell line and reporter spokes identify candidate interactor or near neighbor proteins. Direct protein-protein interactions that have been previously experimentally demonstrated via STRING annotation are represented by additional edges between colored nodes and, in the cases of Sec61β and Ribophorin I, green nodes for members of their native heterooligomeric complexes, which serve as important internal controls. Dark blue nodes and green nodes with asterisks by the gene name are proteins comprising the top GO category indicated underneath each plot and in the included table at the bottom of the figure. For Sec61β (**Figure 7**), a prominent interactor was Sec61*α*, as would be expected if the reporter was assembled into the native Sec61 translocon. The enriched BioID neighborhood set for Sec61β also included membrane biogenesis enzymes, e.g., the stearoyl desaturase SCD, the IP_3_ receptor/calcium channel ITPR3, and the calcium ATPase ATP2B1. GO analysis of the enriched Sec61β BioID interactome set yielded the category “organelle membrane” as a high probability functional gene set. The enriched interactome set for Ribophorin I (RPN1 in figures) (**Figure 7**) includes STT3A, STT3B, and RPN2, (subunits of the OST complex), accessory components of the translocation machinery, such as SSR1, the IP_3_ receptor/calcium channel ITPR2, and the stearoyltransferase SOAT1. GO analysis of the enriched ribophorin I BioID interactome set yielded the category “transport” as a high probability functional gene set. The LRRC59 (**Figure 7**) enriched interactome was particularly interesting as it included ER membrane proteins either predicted or demonstrated to function as RNA binding proteins, including MTDH (AEG-1), RRBP1 (p180), and CKAP4, as well as SRP68 (68 kDa subunit of the signal recognition particle) (Castello et al., 2012, 2016; Hentze, Castello, Schwarzl, & Preiss, 2018). This functional enrichment is consistent with recently published work demonstrating a function for AEG-1 and RRBP1 in RNA anchoring to the ER, and implicate LRRC59 in translational regulation on the ER (Hsu et al. 2018; Cui, Zhang, and Palazzo 2012; Ueno et al. 2012); the GO category “poly(A)RNA-binding” reflects this enrichment. The Sec62 BioID chimera dataset contained very few proteins that met the “enriched” cutoff criteria, with the few that did representing likely false positives (ER lumenal and mitochondrial matrix proteins) (see supplementary proteomic data set)(**Figure 8B**). Also, and whereas we predicted Sec63 in the Sec62 dataset, Sec63 appeared similarly labeled in the LRRC59 and Sec61β datasets and by selection criteria is shared (**Figure 8B**).

**Figure 7.**
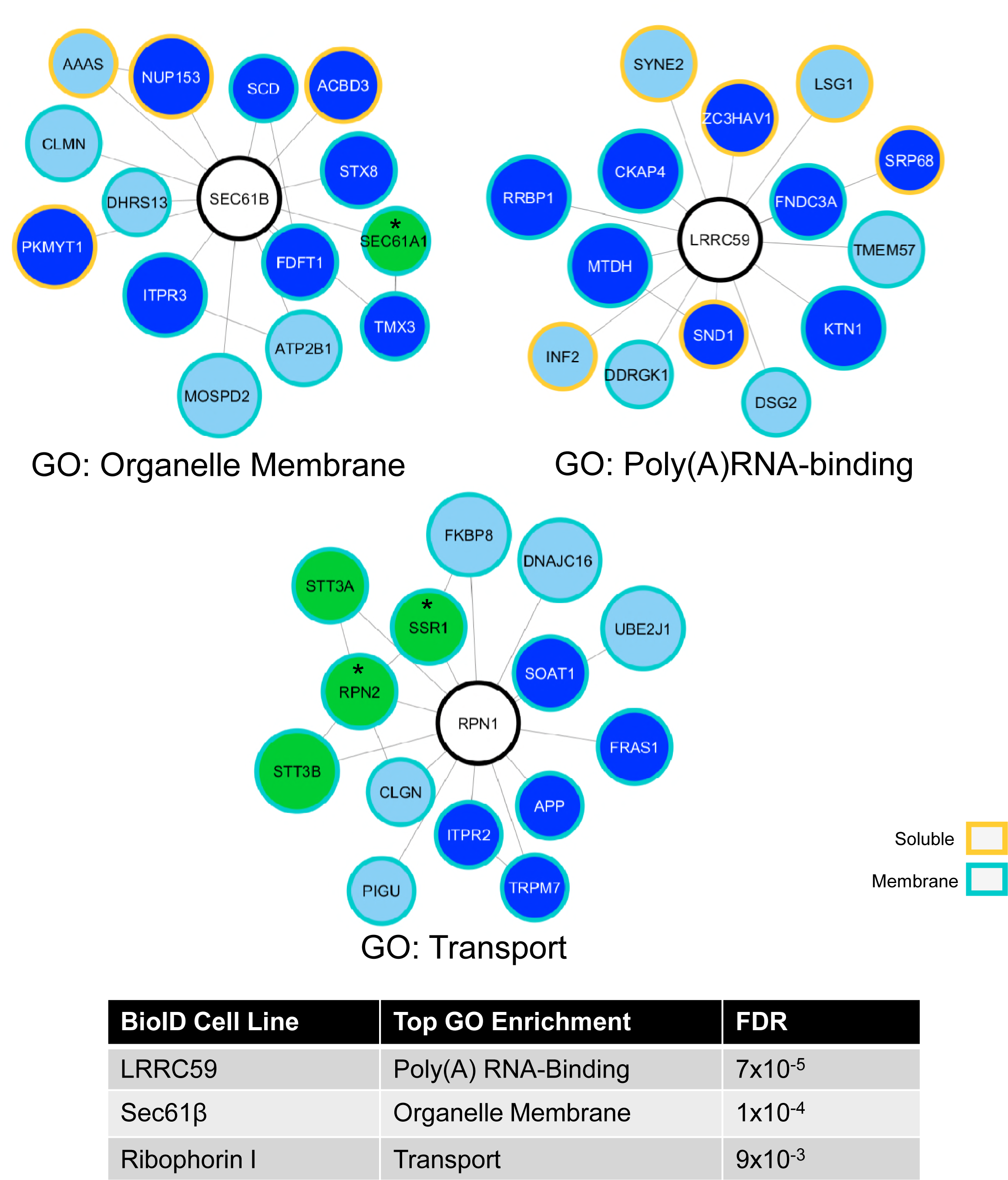
BioID reporters reside in distinct protein neighborhoods. Cytoscape plots of enriched proteins for the Sec61β, LRRC59 and Ribophorin I BioID reporter cell lines reveal different functional enrichments for proximally labeled proteins. Center nodes indicate the chimera protein from each cell line while the surrounding nodes develop from the proteomic datasets. Sizes indicate ranked normalized counts with the largest nodes having the highest values. Green nodes indicate stable, well-characterized protein oligomeric complexes, dark blue nodes are proteins comprising the top GO category indicated underneath each plot and in the appended table. Asterisks denote proteins that are in both established oligomeric complexes and GO categories. Borders indicate whether the protein is a membrane or soluble protein.

**Figure 8.**
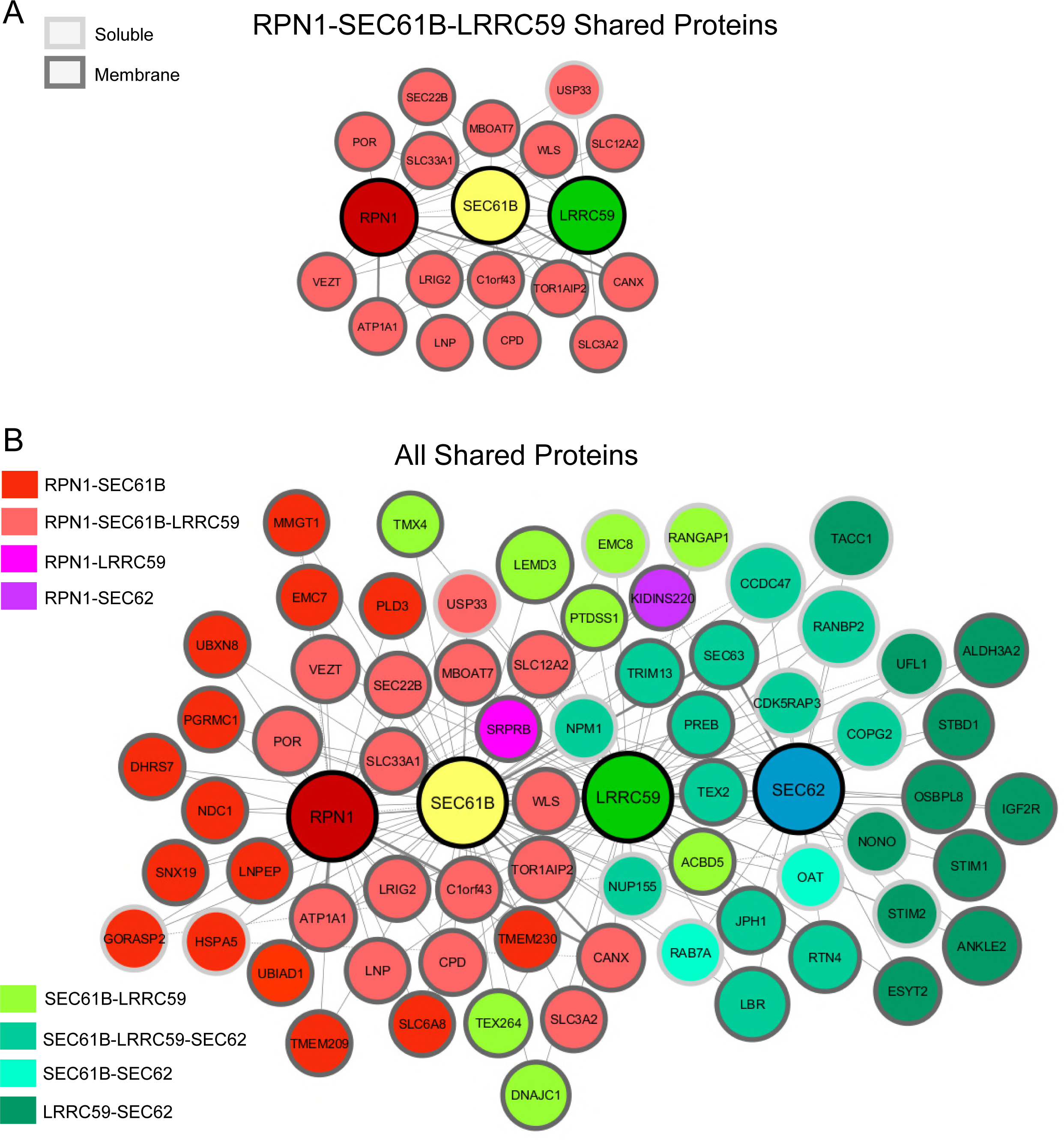
Shared proteins comprise the majority of the proteomic datasets and define common components of mesoscale-ordered ER membrane domains. **A)** The three chimera proteins, Ribophorin I (RPN1), LRRC59 and SEC61β (SEC61B), implicated in ribosome binding, with common, shared proteins. **B)** Cytoscape plot of shared proteins confirms several established protein-protein interactions from experimental evidence (dotted edge lines, bold lines if connected to a reporter node). Size of the nodes are based on highest normalized count from the shared reporters. Proteins shared by specific chimera are distinguished by the indicated color scheme. LRRC59 share more proteins with the SEC61B and RPN1 reporters than SEC62. Files containing all membrane protein raw MS data is contained in Scaffold files as Figure 8 – source data 1 and 2.

The binning scheme (“enriched” and “shared”) is useful for highlighting near-neighbor interactions and their functional enrichments. As noted, however, the proteomics dataset also revealed crosstalk between the BioID chimera and the wild-type candidate ribosome interacting proteins, though these interactions were not above the high stringency cutoff used to define near proximity interactions. The Cytoscape plot illustrated in **Figure 8A** depicts these lower threshold interactions, with the reporter nodes illustrated in yellow, green and dark red, representing Sec61β, LRRC59, and Ribophorin I, respectively, with the remaining “shared” dataset illustrated in **Figure 8B**. In summary, mass spectrometric analysis of the protein neighborhoods/interactomes identified by the BioID method confirm that for two of the candidate ribosome interactors, Sec61β and Ribophorin I, the reporter chimera reside in proximity to their established native oligomeric complexes and intriguingly, three of the chimera reveal distinct protein neighborhoods whose residents are enriched for different ER functions, notably poly(A)RNA-binding (LRRC59), while sharing numerous ER resident proteins functioning in protein biogenesis and other ancillary ER functions (e.g., lipid synthesis and calcium transport).

### Proximity Labeling of ER-Bound Ribosomes by Candidate Ribosome Interactors

In our initial mass spectrometric screens of candidate ribosome interacting proteins, biotinylated ribosomal proteins were largely absent from the datasets, which we subsequently determined reflected inefficient elution of densely biotinylated proteins from the Neutravidin beads. In addition, whereas we had presumed that ribosomal proteins, being highly basic and lysine-rich, would be very receptive to BioID labeling, SDS-PAGE analyses of ER-derived biotinylated ribosomes revealed that only a small subset of ribosomal proteins were targets for BioID labeling (**Figure 9).** As an alternative analytical strategy for analyzing candidate ribosomal proteins labeled via the BioID method, we conducted labeling experiments as above but first enriched for the ribosome fraction by ultracentrifugation and analyzed ribosome BioID patterns separately.

**Figure 9.**
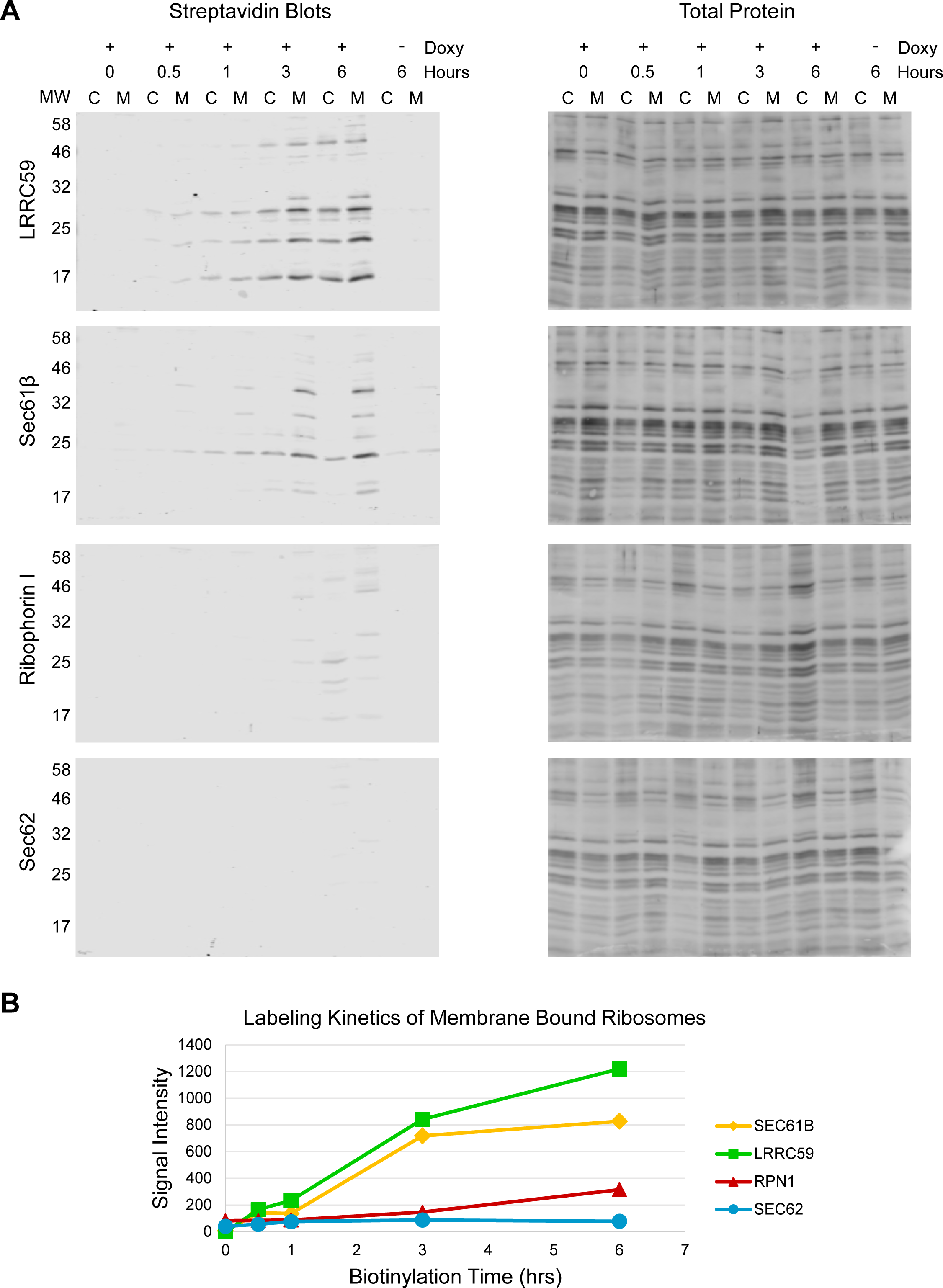
BioID labeling kinetics of cytosolic and ER-bound ribosomes. **A)** Streptavidin blots and related total protein analysis (india ink stains) of the ribosome pellets prepared from samples depicted in Figure 4. **B)** Quantification of the summed lane intensity from membrane-bound ribosome lanes, plotted against time of added biotin using data from the blots shown in panel **A. C** = cytosol, **M** = membrane, **MW** = relative molecular weight in kDa. Data depicted is representative of two biological replicates.

The Sec61β BioID reporter, being a subunit of the Sec61 translocon, was found to efficiently label ER-bound ribosomes, as was the LRRC59 reporter, though itself not a translocon subunit (**Figure 9,** 6 hr time points). Ribosome labeling was not observed at the shorter labeling periods for the Ribophorin I BioID chimera, or at any of the time points examined for the Sec62 BioID chimera (**Figure 9**), though both have been previously reported to bind ribosomes *in vitro* (Harada et al., 2009; Müller et al., 2010). As we report above for the protein interactomes, the restricted ribosome BioID labeling patterns we observed are suggestive of a high degree of spatial organization and stability. Negative data must be interpreted with caution, however. In the case of the Ribophorin I reporter, the ribosome-OST interaction may be too short lived for efficient labeling. Consistent with this interpretation, recent cryoEM studies of ER microsomes have reported two distinct Sec61 translocon environments distinguished by the presence or absence of OST, where it is noted that OST recruitment to the translocon may be transient, being present for the brief interval of N-linked sugar addition (Pfeffer et al., 2015; Wild et al., 2018). In addition and/or alternatively, the Ribophorin I chimera may be compromised in its ability to associate with the Sec61 translocon and thus to report on translocon-bound ribosomes (Braunger et al. 2018).

For the same rationale used in the studies of near-neighbor protein-protein interactions (**Figure 4**), we performed time course studies of ribosome labeling, and examined labeled ribosome distributions in the cytosol and ER compartments (**Figure 9**). As shown, the distinct ribosome labeling patterns seen for the LRRC59 and Sec61β BioID reporters were evident within 0.5 to 1 hour of biotin addition, enriched in the ER-bound ribosome fraction, and with a small fraction of labeled ribosomes recovered in the cytosol. The labeling pattern and relative ratio of ER to cytosolic ribosome labeling, most evident in the LRRC59 BioID reporter line, did not vary substantially over the 6 hour time course of the experiments (**Figure 9**). At present it is not known if the biotin-labeled cytosolic ribosomes represent ribosomes that were labeled in the ER-bound state and subsequently exchanged to the cytosol, or if the BioID chimera labeled both free cytosolic and ER-bound ribosomes. That the ribosomal protein labeling patterns are nearly identical in the ribosomes from both compartments suggests the former. This phenomenon is currently under investigation. As in the experiments illustrated in **Figure 4**, the patterns evident at early time points intensified as a function of labeling time, but did not diversify, suggesting a highly restricted spatial orientation of the BioID reporter-ribosome interface. This phenomenon is further characterized in the analysis depicted in **Figure 9B**, which illustrates the kinetics of the summed signal intensities of the biotinylated ribosomal proteins, for all BioID reporters. Notably, the ribosomal protein labeling kinetics of the LRRC59 and Sec61β BioID reporters are similar, suggesting that the two reporters undergo similar near-neighbor lifetime interactions with membrane-bound ribosomes.

To assess the functional impact of LRRC59 and Sec61β BioID reporter-mediated biotinylation on ribosome function, sucrose gradient sedimentation experiments were performed (**Figure 10A**). The data in **Figure 10B and C** reveal the presence of biotinylated ribosomal proteins in the subunit (**Figure 10B**) and polysome (**Figure 10C**) fractions, where subunit identification was confirmed by sucrose gradient centrifugation and RNA gel analysis of 18S and 28S rRNA distributions, demonstrating that for both reporters, biotin-labeled ribosomes are functionally engaged in translation. The abundant biotin labeling present in the proteins at the top of the gradients represents the ER membrane proteins present in the detergent extracts. These data indicate that LRRC59 and Sec61β BioID-mediated biotin labeling did not compromise ribosome function.

**Figure 10.**
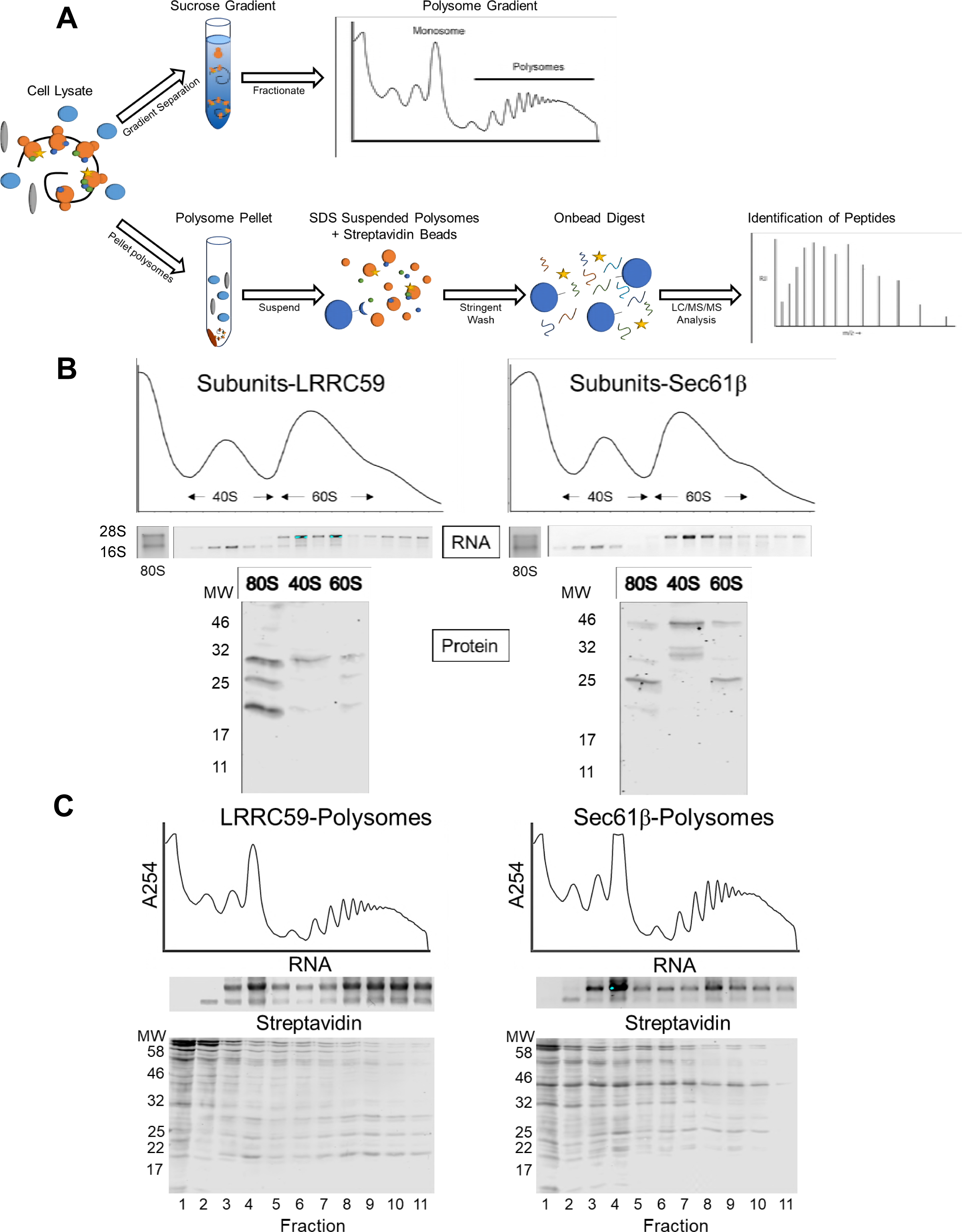
BioID reporter labeling of ER-associated ribosomes in LRRC59 and Sec61β BioID reporter cell lines does not impair translation function. **A)** Experimental schematic illustrating the biotin-tagged ribosome isolation and analysis protocol. **B)** Velocity sedimentation analysis of ribosomal small and large subunits derived from LRRC59 and Sec61β BioID reporter cell lines. Illustrated are the A_254_ traces, RNA gel micrographs depicting 18S and 28S rRNAs. Also illustrated are streptavidin blots of the small and large ribosomal subunits from the reporter cell lines, with 80S ribosomes as comparison. **C)** To determine if BioID-mediated biotinylation altered ribosome function, polyribosomes were fractionated by sucrose gradient velocity sedimentation and biotin-labeled protein distributions analyzed by streptavidin blots of the precipitated gradient fractions. RNA gels of the gradient fractions are included to confirm ribosome migration.

Mass spectrometric analysis of the on-bead digested ribosome fraction revealed a small subset of ribosomal proteins (**Figure 10A, 11**). Consistent with the overall streptavidin labeling patterns of gradient purified ribosomes, determined by streptavidin immunoblot analyses, both BioID reporters labeled a common set of ribosomal proteins: L7a, L14, L23a, and LA2 (not shown) whose locations on the ribosome, illustrated in **Figure 11A** are regionally clustered. Intriguingly, these shared proteins distribute in regions adjacent to the peptidyl transferase and nascent peptide exit site (Wilson & Doudna Cate, 2012). Two ribosomal proteins were highly enriched in only one dataset. RPL17, enriched in the Sec61β dataset, is located near the nascent chain exit site and has been demonstrated to serve important roles in transmembrane domain sensing and signaling to the peptidyl transferase, a function consistent with its appearance in the Sec61β interactome (Lin, Jongsma, Pool, & Johnson, 2011; Zhang, Wölfle, & Rospert, 2013). RPS3A, enriched in the LRRC59 dataset, is located near the mRNA exit site and has been shown to interact with the transcription factor CHOP (Cui et al. 2000). These data are consistent with cryoEM data depicting a specific and spatially constrained interaction between the ribosome and the translocon, of particular relevance to the Sec61β BioID reporter (Voorhees et al., 2014). The LRRC59 BioID reporter, which resides in an ER membrane neighborhood enriched in integral membrane RNA binding proteins, also resides in proximity to bound ribosomes, consistent with a function in coupling translating ribosomes to translocons (Reid & Nicchitta, 2015). These data place LRRC59 in an important ER locale with complementary enrichments in poly(A)RNA binding and translation. The precise role(s) performed by LRRC59 in this environment awaits further study.

**Figure 11.**
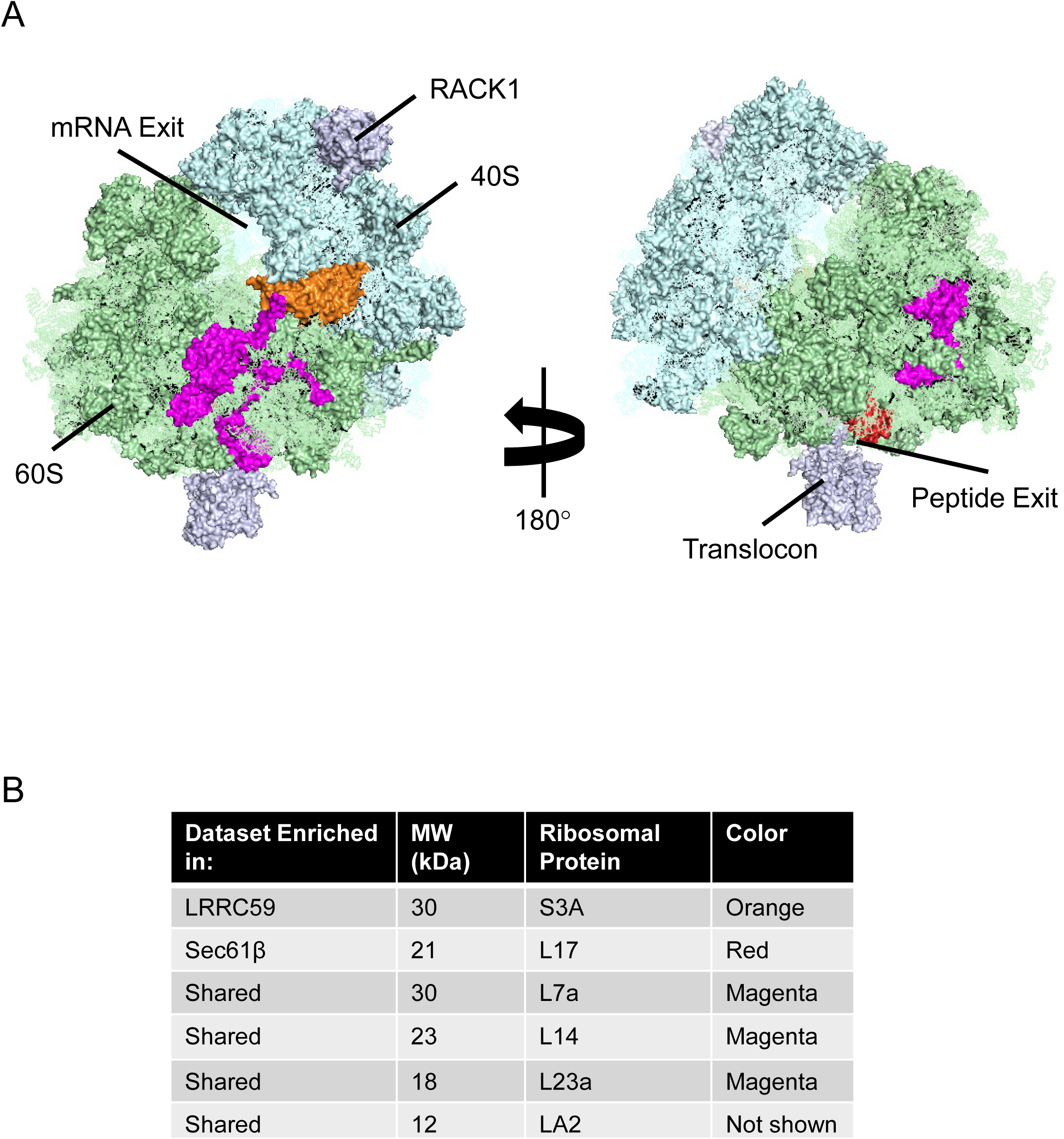
MS analysis of biotin-labeled ribosomal proteins reveals distinct BioID labeling patterns, suggestive of restricted steric interactions with the BioID reporters. **A)** MS/MS identified ribosomal proteins were mapped onto a PDB structure of the ribosome bound to the translocon (PDB: 3J7R). High confidence biotin labeled ribosomal proteins are mapped to the ribosome and several ribosomal features are labeled for orientation. MS experiments were performed in biological duplicate. **B)** Table of the five ribosomal proteins identified from the mass spec datasets with high confidence. Files containing all membrane protein raw MS data is contained in Scaffold files as Figure 11 – source data 1

### Domain-specific RNA-seq reveals regional mRNA enrichments and broad translation functions for ER-bound ribosomes

A primary objective of this study was to examine the translational landscape of the ER, using candidate ribosome interacting proteins as probes for identifying the composition, organization, and translation activities of ribosome-ER association sites. As noted above, we identified robust near-neighbor interactions between ER-bound ribosomes the translocon subunit Sec61β, and the candidate ribosome receptor LRRC59. Intriguingly, the membrane protein neighborhoods for the two bound ribosome interactors displayed divergent functional enrichments, consistent with functions in protein translocation and mRNA translation, respectively. To extend these findings to the translational status of these domains, biotin-tagged, ER-associated ribosomes were purified from the Sec61β and LRRC59 BioID cell lines and the associated mRNAs identified by RNA-seq (**Figure 12**). The experimental methodology is summarized in **Figure 12A**. Following the biotin labeling period, the cytosolic, free ribosome fraction was released via the sequential detergent extraction method noted above and the ER-bound ribosome fraction recovered by detergent solubilization of cytosol-depleted cells. The ribosome fraction was then separated from the co-solubilized membrane proteins by chromatography on Sephacryl S-400 gel filtration media. Biotinylated ribosomes, which are recovered in the S-400 void fractions, were isolated by avidin-magnetic bead capture, the total RNA fraction isolated, and cDNA libraries prepared for deep sequencing. To correct for background mRNA contributions, parallel isolations were performed with empty vector parental cell lines and cDNA libraries from these mock purifications deep sequenced in parallel.

**Figure 12.**
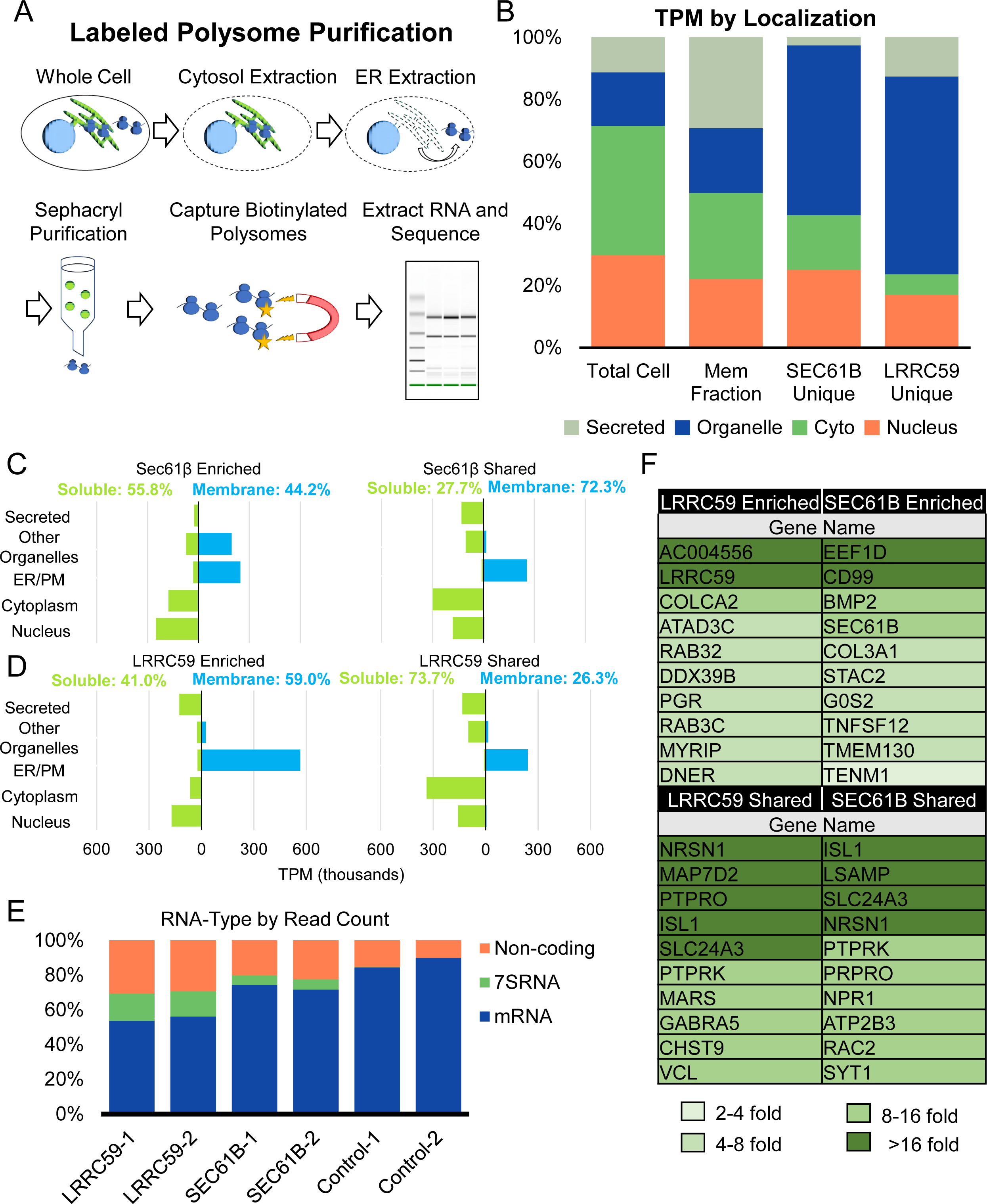
RNA-seq analysis of BioID reporter-labeled polysomes shows divergent transcriptomes and demonstrates that ER ribosomes engage in the translation of cytosolic and secretory protein-encoding RNAs. **A)** Schematic of experimental protocol for capturing biotin-labeled polysomes. **B)** Final subcellular distributions of **proteins** encoded by ribosome-associated RNAs. Stack plots of RNA-seq TPM reveal an enrichment for organellar membrane proteins (DeepLoc1.0) compared to the total mRNA distribution by TPM for membrane and total cell (LocTree3, using datasets from (Reid and Nicchitta, 2012). The transcriptomes from Sec61β and LRRC59 labeled ribosomes diverge from the total and secretory/membrane distributions, and from one another. **C)** TPM analysis of the enriched and shared mRNAs for the Sec61β datasets showing both subcellular distributions and membrane prediction of the encoded proteins. TPM analysis of the enriched and shared mRNAs for the LRRC59 datasets showing both subcellular distributions and membrane prediction of the encoded proteins. Plasma membrane (PM); Transcripts per million (TPM). **E)** Read count analysis of raw counts shown for each of the datasets by percentage of those that aligned to the human genome and counted by htseq-count as described in the methods. **F)** Table of top ten genes by fold change value for enriched and shared datasets color coded by fold enrichment over the control datasets.

In **Figure 12B**, mRNA subcellular category distributions in the shared, Sec61β-enriched, and LRRC59-enriched pools are illustrated. As expected, the ER-associated mRNA transcriptome (**Figure 12B,** Mem Fraction) differs substantially from the total cellular mRNA transcriptome, (**Figure 12B,** Total Cell) showing a substantial enrichment in secretory- and membrane protein-encoding transcripts. As previously reported, the ER-associated mRNA transcriptome contains a substantial representation of cytosolic protein-encoding mRNAs (Reid and Nicchitta 2012; Jan, Williams, and Weissman 2014; Chartron, Hunt, and Frydman 2016). This overall distribution is generally represented, per category, in the shared and Sec61β- and LRRC59-enriched categories, though with significant variations. These differences are further explored in **Figure 12C and D**, which depicts the TPM distributions of mRNA cohorts in the shared and enriched gene sets. As with the total ER-associated transcriptome, when expressed as a relative fraction of the total ER-associated mRNAs, the majority mRNA fraction in the shared category encodes cytosolic proteins (Reid & Nicchitta, 2012). As reported previously, and although relatively abundant and broadly ER-represented, this mRNA cohort is about 2-fold de-enriched relative to the cytosol resident fraction, similar to the fractional distributions reported in yeast and mammalian cell lines (Reid and Nicchitta 2012; Jan, Williams, and Weissman 2014; Chartron, Hunt, and Frydman 2016). In comparison to the total ER-associated and reporter-enriched cohorts, the shared gene set has a somewhat higher representation of cytosolic protein-encoding mRNAs, indicating that in general, this cohort of mRNAs is not selected into either the Sec61β or LRRC59-translation domains. In contrast, membrane protein-encoding mRNAs are substantially enriched in both translation domains (**Figure 12C,D**). Furthermore, individual organelle categories showed unexpected and divergent enrichments. For example, the Sec61β translation domain is enriched for nuclear genes yet de-enriched for secreted genes. Intriguingly, the enriched gene sets for the two translation domains are divergent in mitochondrial genes, with the shared gene set being enriched in matrix (soluble) genes and the enriched sets in genes encoding mitochondrial membrane proteins. As with the cytosolic genes, these data indicate that mitochondrial matrix protein-encoding mRNAs are not selected into either of the translation domains whereas mitochondrial membrane protein-encoding mRNAs are. Furthermore, and quite unexpectedly, GO analysis of the mitochondrial genes in these categories revealed further specification, with the Sec61β translation domains being enriched for mitochondrial outer membrane protein genes and the LRRC59 translation domains being enriched for inner membrane electron transport membrane proteins (**Table 2**). Also displayed in **Tables 1,2** are the highest confidence GO term enrichments for the principal divergent gene sets, which demonstrate that the two examined translation domains display both specification, in the enriched gene sets, and generalization, in the shared gene sets.

**Table 2:** GeneID TPM Enriched Subcellular Components

| Dataset | GO Term | FDR |
| --- | --- | --- |
| LRRC59 – Cell membrane | Cell-cell signaling | 3.37x10<sup>-2</sup> |
| LRRC59 – Extracellular | Extracellular matrix organization | 2.25x10<sup>-4</sup> |
| SEC61B – Nucleus | Negative regulation of transcription | 4.07x10<sup>-2</sup> |
| SEC61B – Cytosol | Neurotrophin signaling pathway | 5.99x10<sup>-4</sup> |
| LRRC59-mitochondria | Electron transport | 4.63x10<sup>-4</sup> |
| SEC61B-mitochondria | Mitochondrial membrane | 6.34x10<sup>-7</sup> |
| Shared – mitochondria | Mitochondrial matrix | 3.27x10<sup>-2</sup> |

**Table 1:** GeneID Unique and Shared

| Dataset | GO Term | FDR |
| --- | --- | --- |
| Shared (306) | System development | 3.31x10<sup>-11</sup> |
|  | Membrane | 3.87x10<sup>-12</sup> |
| LRRC59 Unique (145) | Calcium ion binding | 4.77x10<sup>-2</sup> |
|  | Plasma membrane | 2.03x10<sup>-2</sup> |
| SEC61B Unique (155) | Regulation of signaling | 1.60x10<sup>-3</sup> |
|  | Endoplasmic reticulum | 3.60x10<sup>-4</sup> |

At present, the molecular basis for such enrichments are unknown. Intriguingly, for both translation domains, one of the most enriched genes is the parent reporter gene. Thus, ribosomes engaged in the translation of the reporter reside in proximity to their translation product (**Figure 12F**). Such an intimate association may arise if the reporter parent genes encode or associate with an interacting RNA binding protein. Consistent with this view, Sec61β has been identified as a poly(A) RNA binding protein (Baltz et al., 2012), and LRRC59 resides in an RNA binding protein-enriched domain (**Figure 7**).

In addition to the shared and divergent gene cohorts in coding mRNAs, discussed above, the sequencing data also revealed differences in non-coding mRNAs (**Figure 12E**). For the purposes of this study, we focused on the non-coding 7S RNA of the signal recognition particle (Walter & Blobel, 1982). 7S RNA is enriched over control in the BioID reporter translation domains, with a higher representation in the LRRC59 vs Sec61β translation domains. Consistent with the 7S RNA enrichment in the LRRC59 translation domain, SRP68, the 68kDa SRP protein subunit, was enriched in the LRRC59 proteomics dataset and serves as orthogonal validation for the enrichment of SRP in the two translation domains (**Figure 7 and 12E**). In summary, RNA-seq analyses of the mRNAs undergoing translation on ribosomes proximal to the Sec61β and LRRC59 BioID reporters revealed translational specialization, where specific GO category gene sets were enriched in the two domains, and shared translation functions, where numerous genes were common to the two translation domains. Most noteworthy were divergencies in enrichments for genes encoding mitochondrial matrix or membrane proteins, and “self” genes, where the two translation domains enrich for their respective self RNAs. Combined with the membrane protein proteomic data reported above, these data reveal a higher order meso-organization of the ER, with resident membrane proteins, ribosomes and mRNAs, displaying discrete and complementary enrichments, which is consistent with a model where the ER is comprised of both stable and interacting membrane domains functioning as local translation sites and which can be engaged in the translation of enriched subsets of cytosolic and secretory/membrane protein-encoding RNAs.

## Discussion

Here we report on the translational landscape of the ER from the perspective of candidate ribosome interacting proteins and their protein interactome networks. The rationale for this study is rooted in the growing number of reports demonstrating that cytosolic protein-encoding mRNAs undergo translation on ER-bound ribosomes. Indeed, recent analyses indicate that cytosolic protein-encoding mRNAs can comprise the majority of the translation activity of total ER-bound ribosomes (Reid and Nicchitta 2012; Jan, Williams, and Weissman 2014; Chartron, Hunt, and Frydman 2016). These reports raise a number of fundamental questions regarding mechanisms of RNA localization to the ER, the spatiotemporal regulation of ER-associated translation and in particular, and mechanisms of ribosome association and exchange on the ER *in vivo*, for which it is generally accepted that ribosome exchange on the ER are functionally linked to secretory/membrane protein synthesis (Hsu and Nicchitta 2018). An additional challenge is epistemological, where a dedicated role for the ER in the biogenesis of secretory and membrane proteins is well established, though the question of the exclusivity of this role has been raised for many decades and continues to be debated (Mueckler and Pitot 1981; Diehn et al. 2006; Reid and Nicchitta 2012; Reid and Nicchitta 2015; Jan, Williams, and Weissman 2015). Here we used an unbiased proximity labeling approach, BioID, to investigate the near-neighbor environments of both established and candidate ribosome-interacting ER membrane proteins, including Sec61β, a subunit of the Sec61 translocon, Ribophorin I, a subunit of the OST complex, which resides in proximity to the Sec61 translocon (Harada et al., 2009), and LRRC59 (p34), which displays ribosome binding activity *in vitro* (Tazawa et al., 1991). We draw three primary conclusions from these studies; i) the ribosome interactors examined reside in stable, interactome-ordered ER membrane domains; ii) LRRC59 resides proximal to ER-bound ribosomes and thus likely contributes to the totality of ribosome association on the ER; and iii) the mRNA compositions of ribosomes residing in different membrane domains can be distinguished and comprise both selectively enriched as well as shared transcriptome cohorts. Combined, these data reveal a higher order organization of the ER, which we refer to as mesoscale organization by analogy to current understanding of the domain organization of the plasma membrane (Goyette & Gaus, 2017; Kusumi et al., 2012, 2011), and provide early experimental evidence for a “translation center-based” organization of the ER, where distinct ER domains may function in the coordinated biogenesis of functionally related proteins.

Two largely unexpected observations from this work were the findings that the near-neighbor environments of the different BioID reporter constructs did not diversify as a function of labeling time, and that the direct environments of the BioID reporters were heavily biased to ER membrane proteins. We had initially expected that given the 2-D constraints of the ER membrane, the reporter interactomes would diversify as a function of labeling time to reflect random diffusion in the 2-D plane of the ER membrane. To the contrary, their environments became more densely labeled over labeling time courses of many hours, with only a modest increase in labeling diversity. The remarkable stability of the protein labeling patterns is consistent with a domain model where diverse, low affinity interactions between functionally related proteins enable a mesoscale organization of the membrane. In support of this interpretation, GO analysis of the enriched sets of labeled proteins revealed distinct and functionally related gene categories. Importantly, those BioID reporter chimera that are known to be subunits of oligomeric proteins (Sec61β, Ribophorin I) tagged key subunits, Sec61*α* in the case of Sec61β and both STT3A and STT3B in the case of Ribophorin I, indicating that the chimera assembled into native oligomers and thus reported on the environments of the oligomeric complexes. Although by analogy, extensive studies of plasma membrane architecture have provided strong evidence for mesoscale organization with roles for biophysical contributions from distinct lipid species and interactions with cytoskeletal components as important organizing determinants (Chiantia et al., 2008). Whether lipid species or cytoskeletal components contribute to ER membrane organization remains to be determined, though there is evidence for ceramide/sphingolipid domains in the ER as well as both microtubule and actin cytoskeleton interactions with ER resident proteins (Jagannathan et al., 2014; Ogawa-Goto et al., 2007; Savitz & Meyer, 1997).

Another unexpected observation from this work was the strong biotin labeling bias to the ER membrane over cytosolic proteins. Given the diffusion-based mechanism of BioID labeling, we had expected significant labeling of both cytosolic and membrane proteins. While, the exact reason for this bias remains to be determined, we speculate that it reflects both high local concentrations of reactive sites and high residence lifetimes of ER membrane proteins proximal to the reporters, as contrasted with a soluble protein undergoing three-dimensional aqueous diffusion.

A particularly intriguing observation from these studies is the finding that LRRC59 is near translating ribosomes. Although LRRC59 had been previously reported to function as a ribosome binding protein *in vitro*, a function in ribosome association *in vivo* had not been demonstrated (Ichimura et al., 1993; Ohsumi et al., 1993; Tazawa et al., 1991). Indeed, after a decades long search for the ribosome receptor, which yielded the identification of the Ribophorins, LRRC59, p180, and Sec61, among others, research interest has largely focused on the Sec61 complex as the sole ribosome interacting ER protein (Gorlich et al., 1992; Kalies, Görlich, & Rapoport, 1994; Pfeffer et al., 2015; Voorhees et al., 2014). Indeed, substantial structural data supports this functional assignment, but do not exclude the possibility that additional ER proteins contribute to the totality of ribosome association with the ER (Blau et al., 2005; Müller et al., 2010; Shibatani, David, McCormack, Frueh, & Skach, 2005; Ueno et al., 2012; Wang & Stefanovic, 2014). In support of this conjecture, the BioID interactome for LRRC59 was highly enriched in proteins with candidate or established RNA binding activity, including MTDH (AEG-1), which we recently demonstrated to be an ER RNA binding protein enriched in membrane protein-encoding mRNAs (Hsu et al. 2018), p180, which has also been demonstrated to have a poly(A)RNA binding function, and CKAP4, which was identified as a candidate RNA binding protein in a number of recent studies (Cui, Zhang, and Palazzo 2012; Ueno et al. 2012; Castello et al. 2012). These findings suggest that LRRC59 may have a previously unrecognized role in coupling ER-associated translation to translocon engagement of the translation product. These data fit with a previously proposed model suggesting ribosome interacting proteins might diffuse in the ER membrane to allow nascent chain engagement with unoccupied translocons (Benedix et al., 2010; Johnson & van Waes, 1999; Reid & Nicchitta, 2015). The data in this report demonstrates that LRRC59 resides in proximity to ER-associated ribosomes and engages an interactome enriched in RNA binding proteins suggesting previously unanticipated roles for this protein in translation on the ER.

Since two BioID reporter chimeras distinctly tagged ER membrane-bound ribosomes this provided an opportunity to investigate the transcriptome organization of the ER. As noted, a primary role for the ER in secretory/membrane protein biogenesis is very well established and both past and recent studies examining the subcellular distributions of mRNAs between the cytosol and ER compartments have strongly affirmed this role (Reid & Nicchitta, 2012; Voigt et al., 2017). Although the interpretation of these data has been debated, studies of mRNAs distributions between the cytosol and ER compartments in yeast, by ER localized BirA-AVI tag labeling or by SRP-directed immunoprecipitation, demonstrate that many cytosolic protein-encoding mRNAs display log_2_ cytosol enrichments of < 1-2, and are thus substantially represented on the ER (Jan, Williams, and Weissman 2014; Chartron, Hunt, and Frydman 2016). This mRNA distribution is similar to data reported in mammalian cells (Reid & Nicchitta, 2012; Voigt et al., 2017). In the current study, we examined the associated transcriptomes of biotin-tagged, ER-bound ribosomes. As with earlier studies, we report that although enriched over the cell transcriptome in secretory/membrane protein-encoding mRNAs, ribosomes residing in proximity to both the Sec61β and LRRC59 BioID chimera contained a significant fraction of cytosolic protein-encoding mRNAs and their respective populations of biotin-tagged ribosomes displayed overlapping yet distinct biotin labeling patterns. The RNA populations for the two cell lines displayed both shared and enriched transcripts and, of high interest, the enriched transcript cohorts differed in GO enrichments, with the Sec61βcohort being enriched in mRNAs encoding ER proteins and the LRRC59 cohort being enriched in mRNAs encoding integral plasma membrane proteins. Particularly interesting was the finding that one of the most enriched transcripts for both reporters was the “self mRNA”. These findings support the concept of translation centers on the ER, where mRNAs encoding proteins of related function are coordinately translated in a coherent, localized manner. It remains to be determined how individual mRNAs are targeted to distinct sites or whether mRNAs may be directly recruited to such sites via binding interactions with ER RNA binding proteins such as AEG-1 or by stably associated ribosomes potentially with heterogeneous composition (Hsu et al. 2018; Mauro and Edelman 2002; Wu et al. 2016; Gilbert 2011; Shi et al. 2017).

In summary, we present both proteomic and transcriptomic data supporting the view that translation on the ER, and the ER membrane itself, is subject to mesoscale organization where cohorts of interacting proteins, ribosomes, and mRNAs, are enriched in distinct domain environments. We suggest that such a mechanism may provide for the efficient assembly of functionally related and/or interacting protein complexes. These data also provide additional evidence in support of a transcriptome-wide function for the ER in proteome expression. The remaining questions are many, but given emerging data on the higher order structural organization and translational organization of different regions and compartments of the cell, notably dendrites, mitochondria, stress granules, and P bodies, these data are consistent with higher organization of transcriptome expression and regulation as an evolutionarily conserved cellular strategy (Banani, Lee, Hyman, & Rosen, 2017; English & Voeltz, 2013; Hudder, Nathanson, & Deutscher, 2003; Uezu et al., 2016; Vance, 2014; Youn et al., 2018).

## Materials and Methods

### Generation of BioID Chimera

Plasmids were from the following sources: pCMV-Sport6-RPN1 (Transomic ID: pCS6-BC010839, TransOMIC, Hunstsville, AL), pCMV-Sport6-LRRC59 (Transomic ID: pCS6-BC017168), Sec61β (Transomic ID: pOTB7-BC BC001734), Neo-IRES-GFP-Sec62 (Richard Zimmerman, Saarland University, Homburg, Germany), pEYFP-N1-BirA* (Scott Soderling, Department of Cell Biology, Duke University Medical Center). Gibson assembly master mix (NEB E2611S, Ipswich, MA) was used with the specified amplified fragments using the primers below to generate all constructs with the indicated BirA* tag including a Gly-Ser-Gly-Ser linker between the protein of interest and BirA*. All resulting constructs were cloned into pcDNA5-FRT/TO for downstream generation of HEK293 Flp-In T-REx cell lines (Thermo Fisher Scientific, Waltham, MA). The BioID tags were placed on the terminus facing the cytosol, for Sec62 we chose the C-terminus to avoid disrupting proposed ribosome interactions (Müller et al., 2010). Sequences were confirmed using a CMV-Forward and BGH-Reverse sequencing primers supplied by Eton Biosciences (Research Triangle Park, NC).

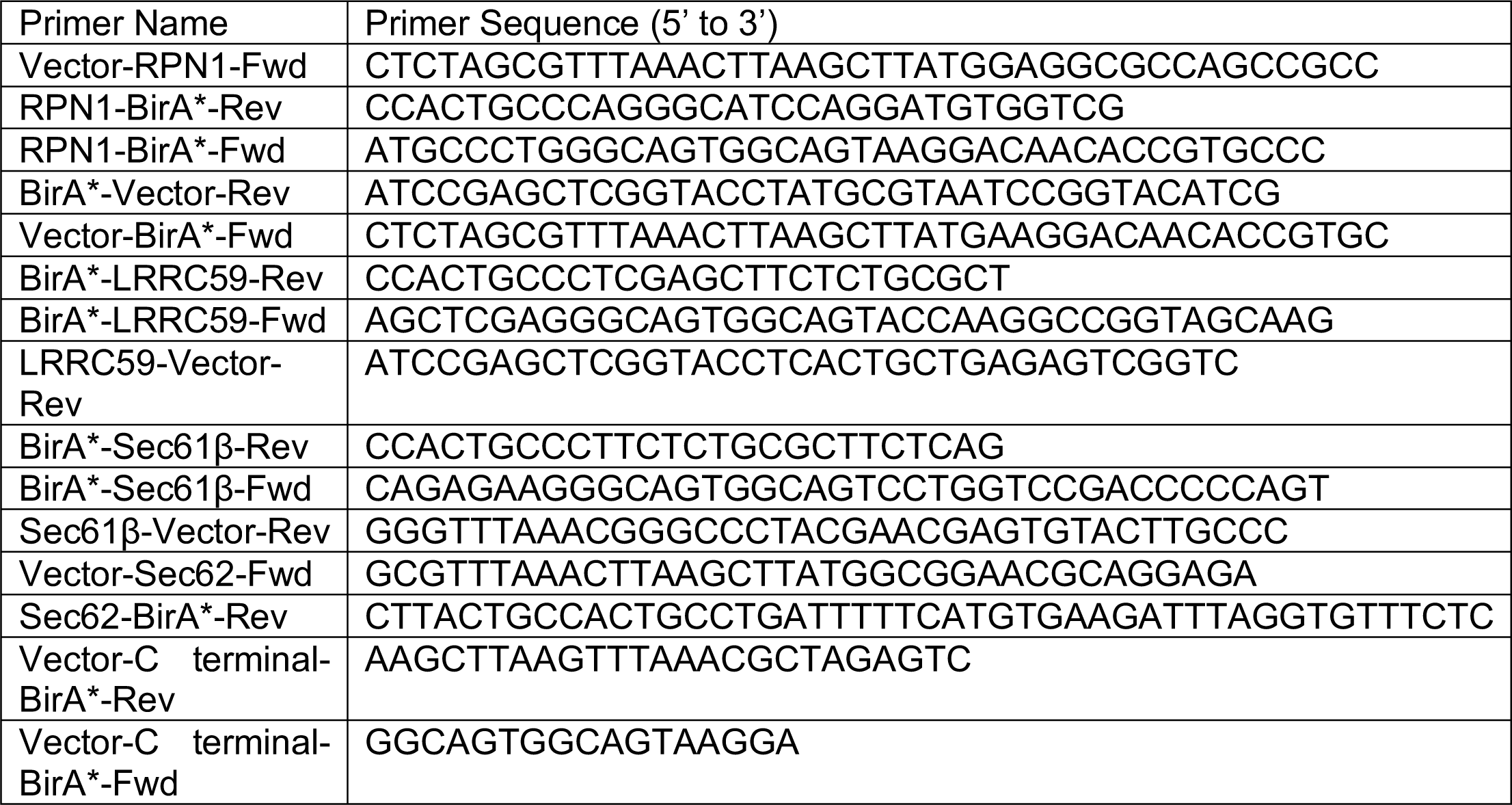

### Generation of HEK293 Flp-In T-Rex cell lines

HEK293 Flp-In T-REx cell lines were generated according to the manufacturer’s instructions (Thermo Fisher Scientific). Cells were transfected in 6-well culture dishes at 80% confluence using 7.5 μL of Lipofectamine 3000 (Thermo Fisher, L3000001) with 0.4 μg of plasmid containing the desired fusion protein and 4 μg of pOGG4 plasmid. Selection with 100 μg/mL hygromycin (MediaTech, 30-240-CR, Manassas, VA) and 15 μg/mL blasticidin (Thermo Fisher, R21001) was started 48 hours after transfection and continued for 2 weeks at which point colonies were identified. A control cell line was generated by recombination of an empty vector pcDNA5-FRT/TO and antibiotic selection for an empty vector matched control.

### Expression of BirA fusion proteins

Expression levels were examined by doxycycline (Sigma Aldrich, D9891, St. Louis, MO) titration and the following doxycycline concentrations were used for each construct: 10ng/mL LRRC59-BioID, 5 ng/mL Sec61β-BioID, 50 ng/mL Ribophorin I-BioID, 100 ng/mL Sec62-BioID. Expression of BioID constructs was performed for at least 16 hr before addition of biotin unless otherwise noted.

### Immunofluorescence Analyses

Cells were plated on poly-lysine coated 22 mm #1.5 coverslips (Globe Scientific, 1404-15, Paramus, NJ). Reporter expression was induced by doxycycline addition and 50 μM biotin added for an overnight labeling. After 16 hours, cells are washed twice with PBS, fixed in 4% paraformaldehyde for 10 min on ice and 10 min at room temperature then washed 3 times with PBS for 5 min each. Cells were permeabilized with a blocking solution of 3% BSA and 0.1% saponin (Sigma Aldrich, S-2149) in PBS for 1 hr at room temperature. Primary staining was performed in the identical solution supplemented with 1:200 BirA antibody (Sino Biological Inc., rabbit IgG, Wayne, PA) or 1:50 TRAPα antibody (Migliaccio, Nicchitta, & Blobel, 1992) (polyclonal, rabbit IgG) at 4°C overnight. Following 5 x 3min washes of 0.1% saponin in PBS, coverslips were incubated with 1:200 goat anti-rabbit IgG AlexaFluor488 (Thermo Fisher, A-11034), 1:1000 streptavidin-Alexafluor647 (Thermo Fisher, S21374) and 1:10000 DAPI (0.5mg/mL stock solution) mixed in blocking solution at room temperature for 45 min in the dark. Coverslips were washed 5X as above, rinsed and mounted in FluorSave hard mount (CalBioChem, 345789, Burlington, MA) and cured at 4°C overnight prior to imaging.

### Fluorescence Imaging

All imaging was performed on a DeltaVision deconvolution microscope (Applied Precision, Issaquah, WA) equipped with 100x NA 1.4 oil immersion objective (UPlanSApo 100XO; Olympus, Tokyo, Japan) and a high-resolution CCD camera (CoolSNAP HQ2; Photometrics, Tucson, AZ). Images were acquired as Z-stacks (at 0.2μm intervals) at identical exposure conditions across the samples for a given protein. The data were deconvolved using the c program (Applied Precision, Mississauga, ON) and processed further on ImageJ-FIJI software to render maximum intensity projections (as required), merge channels and pseudo color the images. Only linear changes were done to the brightness/contrast values of the images, as required and such changes were applied uniformly across all images in a given experiment.

### Sequential Detergent Fractionation and General Cell Lysis

Cells were washed twice with ice-cold PBS containing 50 μg/mL of cycloheximide (CHX) (VWR, 94271, Radnor, PA) for 3 min each wash. Permeabilization buffer (110 mM KOAc, 25 mM HEPES pH 7.2, 2.5 mM Mg(OAc)_2_, 0.03% digitonin (Calbiochem, 3004010), 1 mM DTT, 50 μg/mL CHX, 40U/mL RNAseOUT (Invitrogen, 10777-019, Carlsbad, CA), Protease Inhibitor Complex (PIC) (Sigma Aldrich, P8340)) was added to cells and incubations performed for 5 min at 4°C. The supernatant fraction (cytosol) was collected and cells rinsed with wash buffer (110 mM KOAc, 25 mM HEPES pH 7.2, 2.5 mM Mg(OAc)_2_, 0.004% digitonin, 1 mM DTT, 50 μg/mL CHX, 40U/mL RNAseOUT, Protease Inhibitor Complex (PIC)). Cells were then lysed in NP-40 lysis buffer (400 mM KOAc, 25 mM HEPES pH 7.2, 15 mM Mg(OAc)_2_, 1% NP-40, 0.5% DOC, 1 mM DTT, 50 μg/Ml CHX, 40U/mL RNAseOUT, Protease Inhibitor Complex (PIC)) for 5 min at 4°C. Both cytosolic and membrane fractions were cleared by centrifugation (15,300 x g for 10 min). Total cell lysis was performed in the ER lysis buffer by incubating cells at 4°C for 10 min. Lysates were cleared by centrifugation as above.

### *In Vitro* BirA* Labeling of Microsomes

Canine pancreas rough microsomes (RM) (Walter & Blobel, 1980) were adjusted to a concentration of 4 mg/mL protein in 500 μL of BirA reaction buffer (20 mM Tris pH 8, 5 mM CaCl_2_, 100 mM KCl_2_, 10 mM MgCl_2_, 3 mM ATP, 1.5 mM biotin, 5 mM phosphocreatine (Sigma-Aldrich, P7936-1G) and 5 μg/mL of creatine kinase (Sigma-Aldrich, C3755-3.5KU)). Purified recombinant BirA*-GST fusion protein was added to a concentration of 10 μg/mL. At 0, 1, 3, 6, and 18 hrs, 100 μL of reaction was removed, flash frozen in an ethanol bath and stored at –80°C prior to Western blot analysis.

### Western blotting

Lysate protein concentrations were determined using a Pierce BCA Protein Assay Kit (ThermoFisher, 23225). SDS-PAGE was performed in 12% acrylamide gels containing 0.5% of trichloroethanol. Gels were UV irradiated for 5 min and imaged using an Amersham Imager 600 (GE Life Sciences, Pittsburgh, PA) to verify protein loading. Gels were then equilibrated in Tris-glycine transfer buffer for 5 min and transferred using a Trans Blot SD Semi-Dry Transfer apparatus (Biorad, Hercules, CA). Blots were blocked in PBS, 3% BSA for 1 hr before primary antibody was added at the indicated dilution and incubated for 2 hr at RT or overnight at 4°C. Goat secondary antibodies (Li-Cor, Lincoln, NE) were matched to the species of the primaries used and diluted 1:10,000. Streptavidin was used at a dilution of 1:20,000. Secondary reagents were incubated for 45 min, washed 5x with TBST and imaged on the Odyssey Clx (Li-Cor). Primaries used: BirA (Abcam #14002, polyclonal, chicken IgG), TRAPα (Migliaccio et al., 1992)(polyclonal, rabbit IgG), tubulin (Iowa Hybridoma Bank, E7, monoclonal, mouse IgG, Iowa City, IA), Sec61β (Gift of Ramanujan Hegde, University of Cambridge, polyclonal, rabbit IgG), LRRC59 (Bethyl Labs A305-076A, polyclonal, rabbit IgG, Montgomery, TX), Sec62 (gift from Richard Zimmerman, polyclonal, rabbit IgG), Ribophorin I (Migliaccio et al., 1992)(polyclonal, rabbit IgG), streptavidin-RD680 (Li-Cor, P/N 925-68079).

### RNA Extraction

As adapted from (Chomczynski & Sacchi, 2006), RNA was extracted from 1 volume of lysate using 2 volume of GT buffer to 0.5 volumes of water-saturated phenol, pH 4.5 and incubated for 5 min at RT before adding 0.8 volume of chloroform. Following centrifugation for 15 min at 10,000xg, 4°C for 15 min, the aqueous phase was recovered, and RNA precipitated by addition of 1.2 volumes of isopropanol and 0.15 volume of 3M sodium citrate pH 5.2. Following incubation at −20°C for 1 hr, RNA was recovered by centrifugation at 10,000xg, 4°C for 20 min. RNA pellets were washed in 70% ethanol, dried, and resuspended in TE buffer (10 mM Tris pH 8.0, 1 mM EDTA). RNA concentrations were determined using a NanoDrop ND-1000 Spectrophotometer (Thermo Fisher Scientific). RNA quality was examined by denaturing formaldehyde gel electrophoresis.

### Glycerol Gradients

As adapted from (Nikonov et al., 2002), reporter construct expressing BioID lines were lysed in 1 ml/10cm dish of homogenization buffer (20 mM Tris pH 7.4, 500 mM NaCl, 1.5% digitonin, 1mM MnCl_2_, 1 mM MgCl_2_, 1mM DTT, PIC) for 30 min at 4°C. Lysates were cleared by centrifugation in a TLA 100.2 rotor at 40,000 rpm for 10 min, 4°C (TL-100 Ultracentrifuge, Beckman Coulter, Brea, CA). 850 μL of the supernatant was then loaded onto a 8-30% glycerol gradient and centrifuged in an SW-41 rotor at 35,000 rpm for 15.5 hr, 4°C (L5-50B Ultracentrifuge, Beckman). Gradients were fractionated into 12 fractions using a Teledyne Isco gradient fractionation system and analyzed by immunoblot.

### Polysome Gradients

Cells expressing BioID constructs were lysed in 50 mM HEPES, pH 7.2, 200 mM KOAc, 1 mM DTT, 2% dodecylmaltoside (ChemImpex Intl Inc, 21950, Wood Dale, IL), 5 mM EGTA, PIC, 1mM DTT, 50 μg/mL CHX for 10 min at 4°C. Cell lysates were cleared at 15,300xg for 10 min, 4°C. 0.8 mL of lysate was loaded onto 15-50% sucrose gradients and centrifuged for 3 hours at 35,000 rpm, 4°C (L5-50B Ultracentrifuge, Beckman). Gradients were fractionated into 12 fractions using a Teledyne Isco (Lincoln, NE) gradient fractionation system and analyzed by immunoblot and denaturing RNA gel electrophoresis.

### Biotin Pulldowns

Adapted from (Firat-Karalar & Stearnsx, 2015): Constructs were expressed as above, with biotinylation reactions performed for 3 hours prior to sequential detergent fractionation. The membrane fraction was obtained and volume adjusted to a protein concentration of ca. 1.3 mg/ml and diluted 1:1 with 100 mM NaCl, 50 mM HEPES pH 7.4 to reduce detergent concentrations. Pierce NeutrAvidin Agarose (Thermo Fisher, 29200) resin was blocked for 1 hr with 1% BSA and washed three times in HEPES buffer. Pulldown reactions were performed overnight at 4°C. Beads were washed with the following buffers twice each for 10 min at RT. Buffer 1: 2% SDS in 50 mM HEPES pH7.2 Buffer 2: 0.1% DOC, 1% Triton X-100, 1mM EDTA, 500 mM NaCl, 50 mM HEPES pH7.5 Buffer 3: 0.5% DOC, 0.5% NP-40, 1mM EDTA, 250 mM LiCl, 10 mM Tris pH 7.4. Beads were then suspended in 50 μL of biotin elution buffer, vortexed, and heated for 15 min at 70°C. Supernatant fractions were combined and concentrated to 50 μL in a Savant SpeedVac Concentrator (Thermo Fisher Scientific).

### Ribosome Pulldowns

Cells were washed with PBS and lysed in NP-40 lysis buffer (as above). Lysates were cleared at 15,300 × g for 10 min and the supernatant fraction overlaid onto a 1M sucrose cushion (2:1, load:cushion). Samples were centrifuged at 80,000 rpm for 25 min (TLA 100 rotor in TL-100 ultracentrifuge, Beckman). Ribosome pellets were washed with PBS before being suspended in 50 mM HEPES, pH 7.4, 100 mM NaCl, 1% SDS, 10 mM EDTA, 1 mM DTT, by Dounce homogenization. Ribosome concentration was determined by the A_260_ absorbance and calculated using the extinction coefficient: 5×10^7^/cm*M (Matasova et al., 1991). Equal amounts of ribosomes were used for pulldowns, as above. Binding reactions were performed by end-over-end mixing for 90 minutes at room temperature. Beads were washed as above and suspended in 20 μL of HEPES buffer and submitted to the Duke Proteomics Core (DPMSR) for on-bead digestion.

### Mass Spectrometry

#### On-Resin Trypsin Digestion

The Dynabead complexes in solution were washed three times with 500 µL of 50 mM ammonium bicarbonate (AmBic) (Millipore Sigma, Burlington, MA). Twenty microliters of 1.0% acid labile surfactant (RapiGest, Waters, Milford, MA) in AmBic was added to each sample followed by an additional twenty microliters of AmBic. Samples were subsequently reduced with 10 mM dithiothreitol (DTT, Millipore Sigma) for 30 minutes at 40°C with shaking, and alkylated using 20 mM iodoacetamide (IAM, VWR Scientific) for 30 minutes at room temperature. Digestion was performed using 500 ng sequencing grade trypsin in AmBic (5 μL at 0.1 μg/μL, Promega, Madison, WI), at 37°C overnight with shaking. Peptides were extracted by decanting supernatant into a separate 1.5 mL Eppendorf (Hamburg, Germany) LoBind tube, and washing the resin with 50 μL additional AmBic, which was also combined with digested peptides. The combined extract was acidified to 1% v/v trifloroacetic acid (Thermo Fisher Scientific), heated to 60°C for 2 hours to cleave the RapiGest surfactant, and lyophilized to dryness.

#### Gel Electrophoresis

Samples were transferred to the DPMSR for one dimensional sodium dodecyl sulfate polyacrylamide gel electrophoresis (SDS-PAGE). 25 μL of sample was combined with 5 μL of 100 mM DTT and 10 μL of NuPAGE^™^ (Thermo Fisher Scientific) 4X loading buffer and samples were then heated to 70°C for ten minutes with shaking. SDS-PAGE separation was performed using 1.5 mm 4-12% Bis-Tris pre-cast polyacrylamide gels (Novex, Thermo Fisher Scientific), 1X MES SDS NuPAGE^™^ Running Buffer (Thermo Fisher Scientific) including NuPAGE^™^ antioxidant. SDS-PAGE separation was performed at a constant 200V for five minutes, gels fixed for 10 minutes, stained for 3 hours, and destained overnight following manufacturer instructions.

#### Gel Band Isolation and Trypsin Digestion

Gel bands of interest were isolated using a sterile scalpel transferred to protein LoBind tubes (Eppendorf) and minced. Gel pieces were washed with 500 μL of 40% LCMS grade acetonitrile (MeCN, Thermo Fisher Scientific) in AmBic, with shaking at 30°C. Gel pieces were shrunk with LCMS grade MeCN, the solution discarded, and the gel pieces dried at 50°C for 3 min. Reduction of disulfides was performed using 100 μL of 10 mM DTT at 80°C for 30 min with shaking, followed by alkylation with 100 μL of 55 mM IAM at RT for 20 min. This liquid was aspirated from the samples and discarded, and gel pieces were washed twice with 500 uL AmBic, and these washes were also discarded. LCMS grade MeCN was added to shrink the gel pieces in each sample, then samples were swelled in AmBic and this process was repeated a second time, finally the gel pieces were shrunk a final time by adding 200 μL of LCMS grade MeCN, and heating for 3 min at 50°C to promote evaporation. Trypsin digestion was performed with addition of 30 μL of 10 ng/ μL sequencing grade trypsin (Promega, Madison, WI) in AmBic followed by 30 μL of additional AmBic. The samples were incubated overnight at 37°C with shaking at 750 rpm. Finally after overnight digestion 60 μL of 1/2/97 v/v/v TFA/MeCN/water was added to each sample and incubated for 30 min at RT and 750 rpm to extract peptides, and the combined supernatant was transferred to an autosampler vial (Waters). Gel pieces were shrunk in 50 μL additional MeCN for 10 min to extract the maximum amount of peptides, which was combined with the previous supernatant. The samples were dried in the Vacufuge (Eppendorf) and stored at −80°C until ready for LC-MS/MS analysis.

#### Qualitative Analysis of On-Resin and Gel Electrophoresis Samples

All on-resin and gel band samples were resuspended in 20 μL of 1/2/97 v/v/v TFA/MeCN/water. The samples were analyzed by nanoLC-MS/MS using a Waters nanoAcquity LC interfaced to a Thermo Q-Exactive Plus via a nanoelectrospray ionization source. 2 μL of each on-resin sample, and 1 μL of each gel band sample was injected for analysis. Each sample was first trapped on a Symmetry C18, 300 μm x 180 mm trapping column (5 μl/min at 99.9/0.1 v/v H2O/MeCN for 5 min), after which the analytical separation was performed using a 1.7 μm ACQUITY HSS T3 C18 75 μm x 250 mm column (Waters). The peptides were eluted using a 90 min gradient of 5-40% MeCN with 0.1% formic acid at a flow rate of 400 nl/min with a column temperature of 55 °C.

Data collection on the Q Exactive Plus mass spectrometer was performed with data dependent acquisition (DDA) MS/MS, using a 70,000 resolution precursor ion (MS1) scan followed by MS/MS (MS2) of the top 10 most abundant ions at 17,500 resolution. MS1 was performed using an automatic gain control (AGC) target of 1e6 ions and maximum ion injection (max IT) time of 60 msec. MS2 used AGC target of 5e4 ions, 60 ms max IT time, 2.0 m/z isolation window, 27 V normalized collision energy, and 20 s dynamic exclusion. The total analysis cycle time for each sample injection was approximately 2 h. The sample run order was chosen to minimize potential carryover and is detailed as follows for the on-resin and gel band samples, respectively: 125-EV, 125-LR59, 125-S61, 1210-EV, 1210-LR59, 1210-S61, EV, LRRC59, SEC62, SEC61B, and RPN1.

#### Database searching

Proteome Discoverer (Thermo Fisher Scientific) was used to generate mgf files from the DDA analyses and the data was searched using Mascot v 2.5 (Matrix Science) with a custom database containing the human proteome downloaded from UniProt combined with common proteins found in BirA experiments and common contaminants. The data was searched using trypsin enzyme cleavage rules and a maximum of 4 missed cleavages, fixed modification carbamidomethylated cysteine, variable modifications biotinylated lysine, deamidated asparagine and glutamic acid and oxidated methionine. The peptide mass tolerance was set to +/-5 ppm and the fragment mass tolerance was set to +/-0.02 Da. False discovery rate control for peptide and protein identifications was performed using Scaffold v4 (Proteome Software, Inc).

### Analysis of Scaffold data

Method adapted from Ritchie, Cylinder, Platt, & Barklis, 2015. For the membrane protein data sets of each biological replicate, hits with 1% FDR at the protein level, 50% peptide match with a minimum of 2 peptides and 2 spectral counts were used for subsequent analysis. Each dataset is first normalized by summing spectral counts for the natively biotinylated proteins-acetyl-CoA carboxylase, propionyl CoA carboxylase, pyruvate carboxylase, and methyl crotonyl-CoA carboxylase subunits – and dividing all spectral counts by this number. Proteins less than 2.5-fold above the same proteins in the respective control dataset were removed. The remaining protein spectral counts for each dataset were averaged and normalized by dividing by the BirA protein spectral counts to account for any differences in reporter expression. Analyses were performed so that any proteins with average normalized counts higher than 2-fold above the same protein in the three other datasets was assigned to the specific cell line as “enriched.” Remaining proteins were analyzed by covariance of normalized counts with a cut-off of 40.0. These proteins were shared between at least two of the cell lines with higher than 2-fold normalized counts of the lowest count. For figure clarity, the Cytoscape plot in **Figure 8B** displays those shared proteins with a covariance of 50.0 or above. For localization prediction, a FASTA file containing the protein sequences was generated and processed on the TMHMM Server v2.0 (DTU Bioinformatics) to identify membrane vs soluble proteins. Localization by organelle (**Fig 6B**) was determined by running the datasets through DeepLoc v1.0 (DTU Bioinformatics) using the Profiles algorithm.

For ribosomal protein data sets, spectral counts were retrieved for only ribosomal protein hits with 90% protein identity, 50% peptide identity with at least 2 peptides. Each experiment dataset was divided by the control, and those exceeding a 2-fold difference were further analyzed. For each candidate, sample spectral counts were divided by the control and proteins with greater than 2-fold difference are termed “enriched” and those below the cutoff are termed “shared”. Those proteins with the same term between the two datasets are kept and mapped onto PDB file 37JR, of the translating ribosome on the translocon.

### Biotinylated Polysome Isolation and Sequencing

Ribosomes were purified from the membrane fractions of sequential detergent fractionation of the indicated BioID cell lines by gel filtration chromatography, collecting the fraction of a Sephacryl S400 column operating at a flow rate of 0.7 mL/min. Dynabeads M-270 Streptavidin beads (ThermoFisher, 65305) and 0.05% Triton X-100 are added to each sample and incubated overnight at 4°C. Beads were washed three times for 10 min at 4°C in high-salt wash buffer followed by suspension in low-salt buffer and extraction of bound RNA using an RNAEasy Kit (Qiagen, 74104, Hilden, Germany). RNA was quantified by Bioanalyzer 2100 analysis (Agilent, Santa Clara, CA) and like samples combined to provide 10 ng of total RNA total. RNA samples were concentrated to 12 μL using E.Z.N.A. MicroElute RNA Cleanup Kit (Omega Bio-Tek, R6247, Norcross, GA) and libraries constructed using Ultra II RNA Library Kit (NEB, E7645S) for biological duplicates.

### Illumina Hi-Seq

Libraries were submitted to the Duke Sequencing and Genomic Technologies for sequencing. Concentration of each library was estimated using Qubit assay and run on an Agilent Bioanalyzer for library size estimation. Libraries were then pooled into equimolar concentration. Final pool was clustered on a HiSeq 4000 Single-Read flow cell. Sequencing was done at 50bp Single-Read. Bcls files generated by the sequencer were then converted into fastq files using Illumina bcl2fastq v2.20.0.422 and reads demultiplexed using the molecular indexes incorporated during library preparation.

### Sequencing Analysis

FASTA files were adapter trimmed using Trimmomatic v0.32 (Bolger, Lohse, & Usadel, 2014), aligned to the human genome, build GRCh38/h38, using HISAT2.0.5 default options for unpaired reads (D. Kim, Langmead, & Salzberg, 2015). Aligned read files were then counted using htseq-count v0.5.4p3 (Anders, Pyl, & Huber, 2015) using options for non-stranded reads, intersection-strict mode, and ‘exon’ as the feature to be counted using a UCSC hg38 GTF annotation file. This GTF file with unique gene IDs and transcript IDs was generated to a genePred file for hg38 using the genePredtoGTF script from kentUtils. Data sets from the two cell lines were analyzed for differential expression versus the control experiments using DESeq2v1.18.1 (Love, Huber, & Anders, 2014). Gene lists were generated by taking the subset with greater than or equal to 2-fold change over the control data set with an adjusted p-value of 0.05 (Benjamini & Hochberg, 1995). Genes coding for protein products were selected for interaction and GO analysis using the STRING database. Localization predication analysis was performed using the DeepLoc1.0 Profiles algorithm (Almagro Armenteros et al., 2017). Transcript per million (TPM) analysis was performed by first calculating reads per kilobase (RPK), summing the RPK values and dividing by 1 million to use as the scaling factor (SF). Individual RPK values were divided by the SF to obtain a gene specific TPM value for the given subset of data for better comparison of the datasets.

## Acknowledgements

We thank the Duke University School of Medicine for the use of the Proteomics and Metabolomics Shared Resource which provided the proteomics service for this study. In particular, we would like to acknowledge Will Thompson, PhD and Sarah Raines of the Duke Proteomics Core facility for very helpful guidance and advice on experimental design and data analysis. We acknowledge key instrumental resources (QE Plus) supported by NIH S10OD012266-01A1. We also thank the Duke Sequencing and Genomic Technologies Shared Resource, in particular Nicolas Devos, PhD, for advice, guidance, and generous support of the deep sequencing studies. We especially thank members of the Nicchitta lab, in particular Jessica Childs, Heather Vincent, and Jason Arne for insightful discussions throughout this project. The E7 monoclonal antibody was obtained from the Developmental Studies Hybridoma Bank, created by the NICHD of the NIH and maintained at The University of Iowa, Department of Biology, Iowa City, IA 52242. This research was supported by funding from the NIH (GM101533-A1, GM118630-A1, CVN).

## SOURCE DATA FILES

### REVIEWER ACCESS TO GEO ARCHIVED FASTQ FILES

*A secure token has been created to allow review of record GSE118873 while it remains in private status:*

**Figure 8-linked proteomics:** Files containing all membrane protein raw MS data are provided as Scaffold files, file name: Figure 8 – source data 1 and 2.

**Figure 11-linked proteomics:** File containing all ribosomal protein raw MS data is contained in a Scaffold file, file name: Figure 11 – source data 1

**Figure 3 – figure supplement 1.**
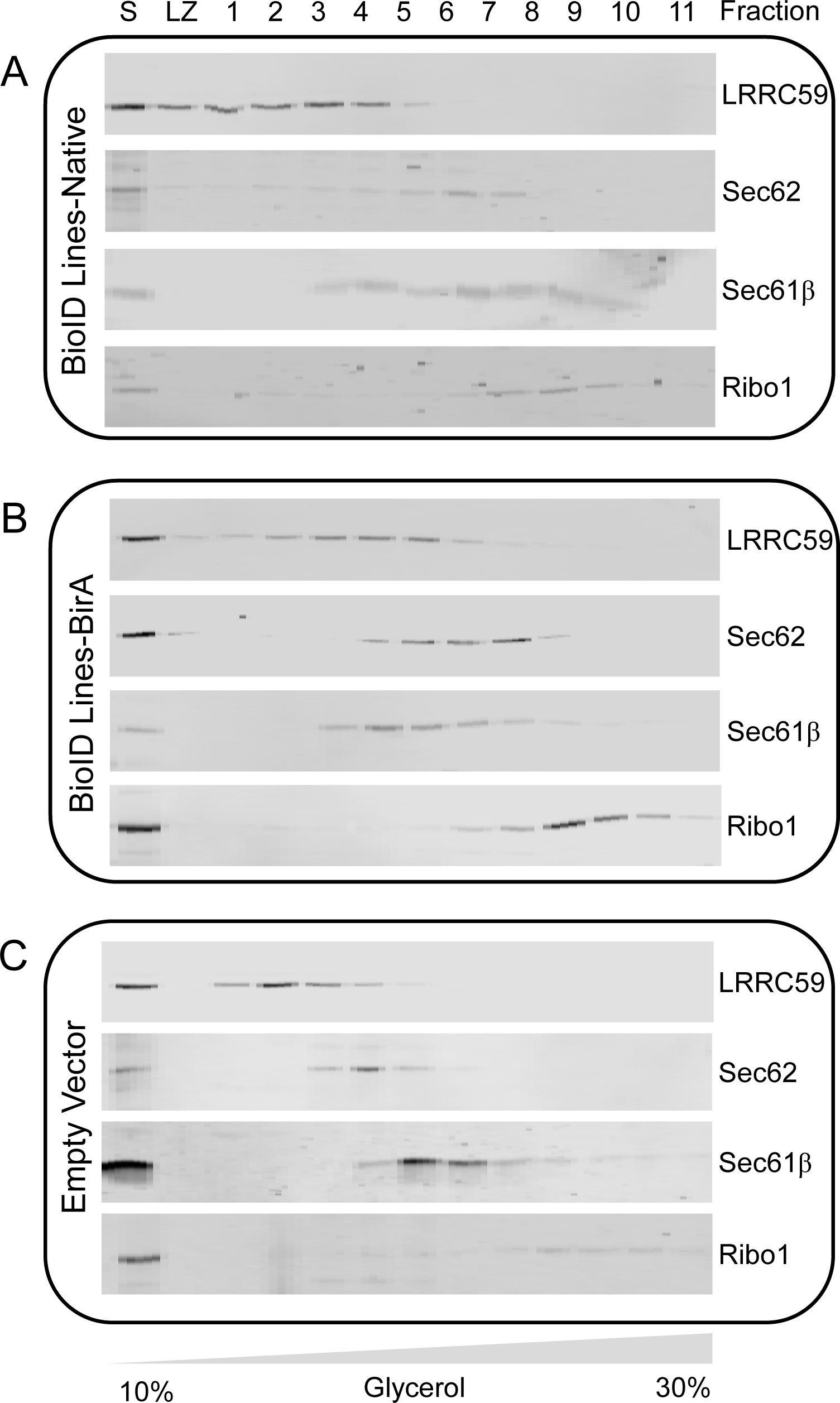
BioID constructs display similar hydrodynamic behavior to native complexes in glycerol gradient sedimentation analyses. **A)** Immunoblot analysis of BioID reporter construct migration in glycerol gradient velocity sedimentation experiments, comparing migration behavior of native and BioID chimera for each BirA*-chimera cell line. Analysis of native proteins. **B)** As in **(A)** but using BirA antisera to compare migration patterns of the BioID fusion proteins with the native protein. As in **(A)** but using an empty-vector control cell line, to compare migration patterns of native proteins as in panels **A** and **B**.

**Figure 12 – figure supplement 1.**
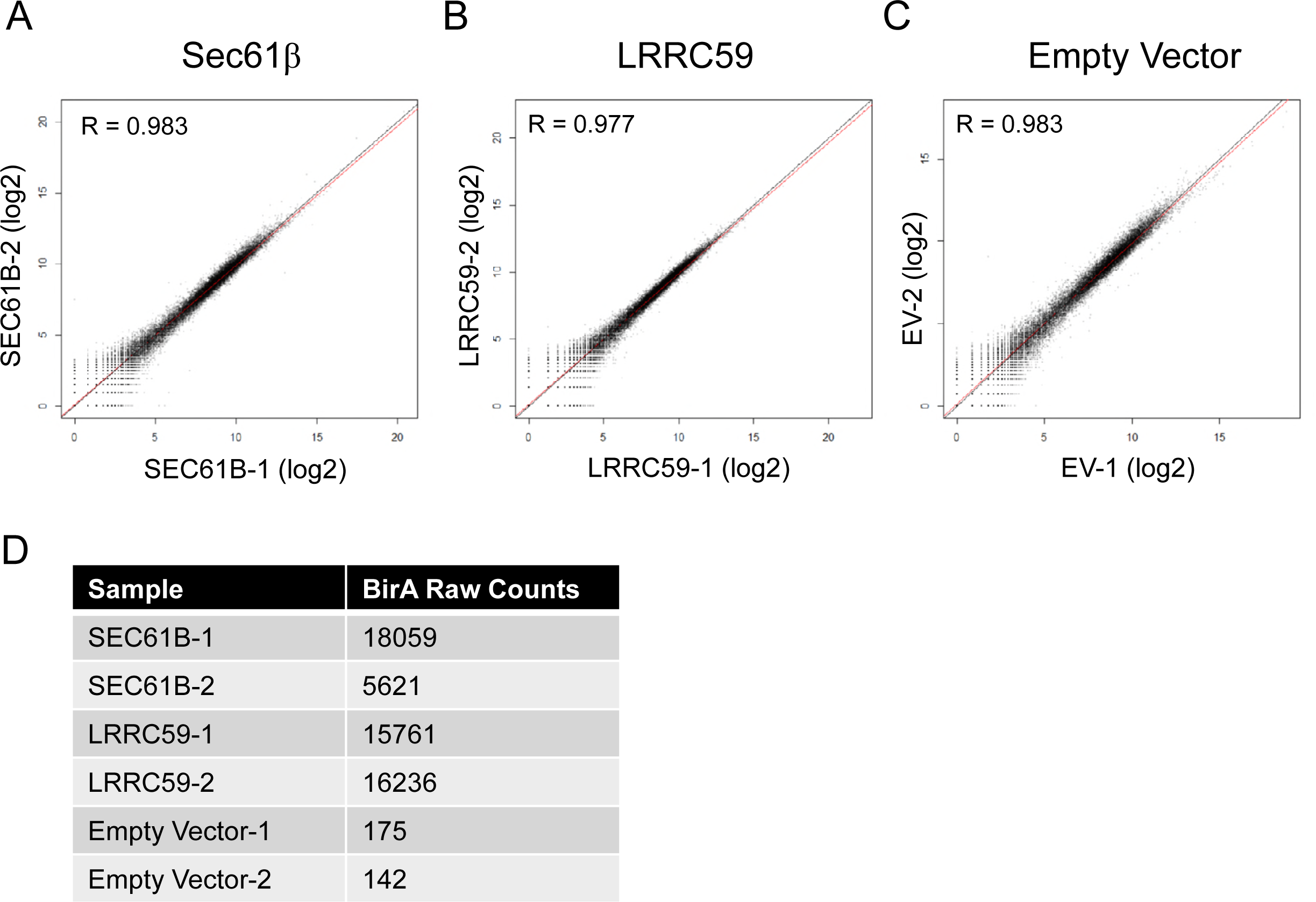
Quality control checks for replicates in RNA-seq libraries. **(A-C)** log_2_ transformed counts from RNA-Seq datasets of biological replicates in the analysis are plotted, revealing high similarity between biological duplicated. Red lines represent regression lines plotted to reveal variation from the midline (black). Pearson correlation coefficient at 95% confidence is shown in upper left part of the graph.**(D)** Raw counts of datasets aligned to the humanized BirA construct sequence used for the BioID chimera demonstrating expression of the BioID chimera and background mapping frequencies in the empty vector lines.

